# Extrachromosomal DNA micronucleation constrains tumour fitness and improves patient survival

**DOI:** 10.1101/2025.04.15.648906

**Authors:** Lotte Brückner, Robin Xu, Jun Tang, Aditi Gnanasekar, Alexander Herrmann, Ivy Tsz-Lo Wong, Shu Zhang, Fengyu Tu, Madison Pilon, Alexander Kukalev, Katharina Pardon, Olga Sidorova, Joshua Atta, Qinghao Yu, Giulia Montouri, Davide Pradella, Mila Ilić, Kentaro Suina, Marco Novais-Cruz, Sarah Kaltenbach, Denise Treue, Mădălina Giurgiu-Kraljič, Sergej Herzog, Ann-Sophie Hollinger, Nadine Wittstruck, Martina Fernandez, Nicole Wolozyn, Finnja Becker, Varvara-Rigina Louma, Xiaowei Yan, Rachel Schmargon, Jan Dörr, David Gamlin, Annika Lehmann, Dennis Gürgen, Matthias Richter, Frank Dubois, Fabrizio Simeoni, Betheney R. Pennycook, Alastair Hamilton, Ralph K. Lindemann, Courtney Dawes, Gabriel Balmus, Barbara Hero, Matthias Fischer, Vineet Bafna, Roel Verhaak, Geoffrey Wahl, Yang Liu, Richard P. Koche, Stamatis Papathanasiou, René Medema, Bastiaan Spanjaard, Andrea Ventura, Ana Pombo, Weini Huang, Monika Scheer, Benjamin Werner, Howard Y. Chang, Paul S. Mischel, Anton G. Henssen

## Abstract

Extrachromosomal DNA (ecDNA) contributes to cancer genome instability by enabling high-copy oncogene amplification, intratumoural heterogeneity and rapid genetic change. Micronuclei (MN) are frequently observed in chromosomally unstable cancers, yet their origins and relevance in ecDNA-driven tumours remain incompletely understood. Here, we investigate the relationship between ecDNA segregation errors and MN formation. We find that ecDNA frequently localizes to MN and represents a prominent source of MN content in ecDNA-positive cancer cells. Mitotic clustering of oncogene-bearing ecDNAs is associated with asymmetric inheritance and mis-segregation into MN. Transfer of oncogenes from ecDNA into MN is accompanied by reduced transcriptional output. Single-MN sequencing shows that individual MN are enriched for ecDNA to a degree that exceeds expectations from stochastic mis-segregation and that in MN oncogenes originating from multiple distinct ecDNAs coalesce. Using live-cell imaging, we observe that cells inheriting ecDNA-positive MN show limited proliferative capacity and an increased likelihood of cell death. In neuroblastoma patients with *MYCN*-amplified ecDNA, higher frequencies of ecDNA-positive MN at diagnosis are associated with improved event-free and overall survival. Together, these findings link ecDNA mis-segregation to MN formation and reduced cellular fitness, suggesting that ecDNA-positive MN may reflect a state of impaired oncogenic ecDNA function with potential relevance for clinical outcome.

## Introduction

Extrachromosomal DNA (ecDNA) has emerged as a major form of cancer genome instability, driving high-copy oncogene amplification, intratumoural heterogeneity, rapid tumour evolution, and therapy resistance^1,2^. Across diverse cancer types, the presence of ecDNA (ecDNA⁺) correlates with poor outcomes compared to tumours with linear chromosomal amplifications or other types of genome instability^3^. EcDNAs are relatively large (approximately 1-2Mb)^4^, chromatinized, circular DNA structures that are enriched for oncogenes, as well as regulatory and immunomodulatory elements^5–7^. They are observed specifically in cancer cells and are distinct from extrachromosomal circular DNA (eccDNA), which comprise smaller, typically non-clonal circular DNA elements that rarely encompass full genes and can also be detected in normal cells^8–10^. Their absence of centromeres on ecDNA^11,12^, permits uneven segregation during mitosis, fuelling heterogeneity and accelerating tumour evolution through non-Mendelian inheritance^11^. Chromosomal tethering mechanisms are thought to promote ecDNA transmission during cell division^13,14^.

Micronuclei (MN) are small, extranuclear bodies initially enclosed by nuclear envelopes containing mis-segregated chromosomes or chromatin fragments arising during mitosis^15,16^, distinguishing them from other extranuclear structures such as exclusomes^17^. They are a hallmark of chromosomal instability. Chromatin in MN can undergo chromothripsis^18–20^ and epigenetic remodelling^21–23^, resulting in long-term genomic and transcriptional alterations. MN can serve as intermediates for ecDNA generation^20^, suggesting reciprocal links between ecDNA and MN biology. Treatment of ecDNA-bearing cells with hydroxyurea or other DNA-damaging agents increases MN frequency and decreases oncogene levels, implying that damaged ecDNA may seed MN formation^24–26^. However, the mechanisms of MN generation in ecDNA⁺ tumours, their molecular composition, and functional consequences remain unknown.

Using cell lines, engineered mouse models, patient tumour samples, single-MN sequencing, and live-cell imaging, we set out to define how micronucleation arises from ecDNA and how this process shapes cancer cell fitness and patient outcome.

## Main

### ecDNA gives rise to oncogene-containing micronuclei

To assess how frequently ecDNA is incorporated into micronuclei (MN) and how faithfully it is retained during cell division, we compared near-isogenic cancer cell lines carrying oncogene amplifications either on ecDNA or re-integrated as homogeneously staining regions (HSRs). (Fig. 1a, Extended Fig. 1). Across three such pairs, COLO 320DM/HSR (*MYC*), GBM39EC/HSR (*EGFR*vIII), and PC3-DM/HSR (*MYC*), ecDNA⁺ cells displayed significantly higher frequencies of oncogene-positive MN (oncogene^+^ MN) than their HSR counterparts (Fig. 1b). In contrast, the frequency of oncogene⁻ MN was similar, indicating that ecDNA mis-segregation contributes to elevated MN frequency (Fig. 1b-c). Lamin B1 expression was higher in oncogene⁺ MN than oncogene^-^ MN in ecDNA⁺ cell lines, suggesting that oncogene+ MN have an intact inner nuclear membrane (Extended Fig. 2a-c). Analysis of 275 patient tumour biopsies from diverse cancer types with FISH-defined amplification status of distinct oncogenes (Fig. 1d, Supplementary table 1) confirmed this pattern: ecDNA⁺ tumours contained a significantly greater fraction of cells with MN than non-amplified tumours or tumours with chromosomal amplifications, and the proportion of oncogene⁺ MN was correspondingly higher (Fig. 1e-f, Extended Fig. 2d). Micronuclei in ecDNA⁺ cells were smaller than those in HSR cells, consistent with their predominant composition of ecDNA rather than larger chromosomal fragments (Extended Fig. 2e). MN abundance positively scaled with overall ecDNA copy number in most tumour types (Fig. 1g–j, Extended Fig. 2f). A binomial segregation model estimated an ecDNA mis-segregation rate of 1.7% (range 0–7.8%), indicating overall efficient, but occasionally faulty, retention during mitosis (Extended Data Fig. 2g).

**Fig 1.**
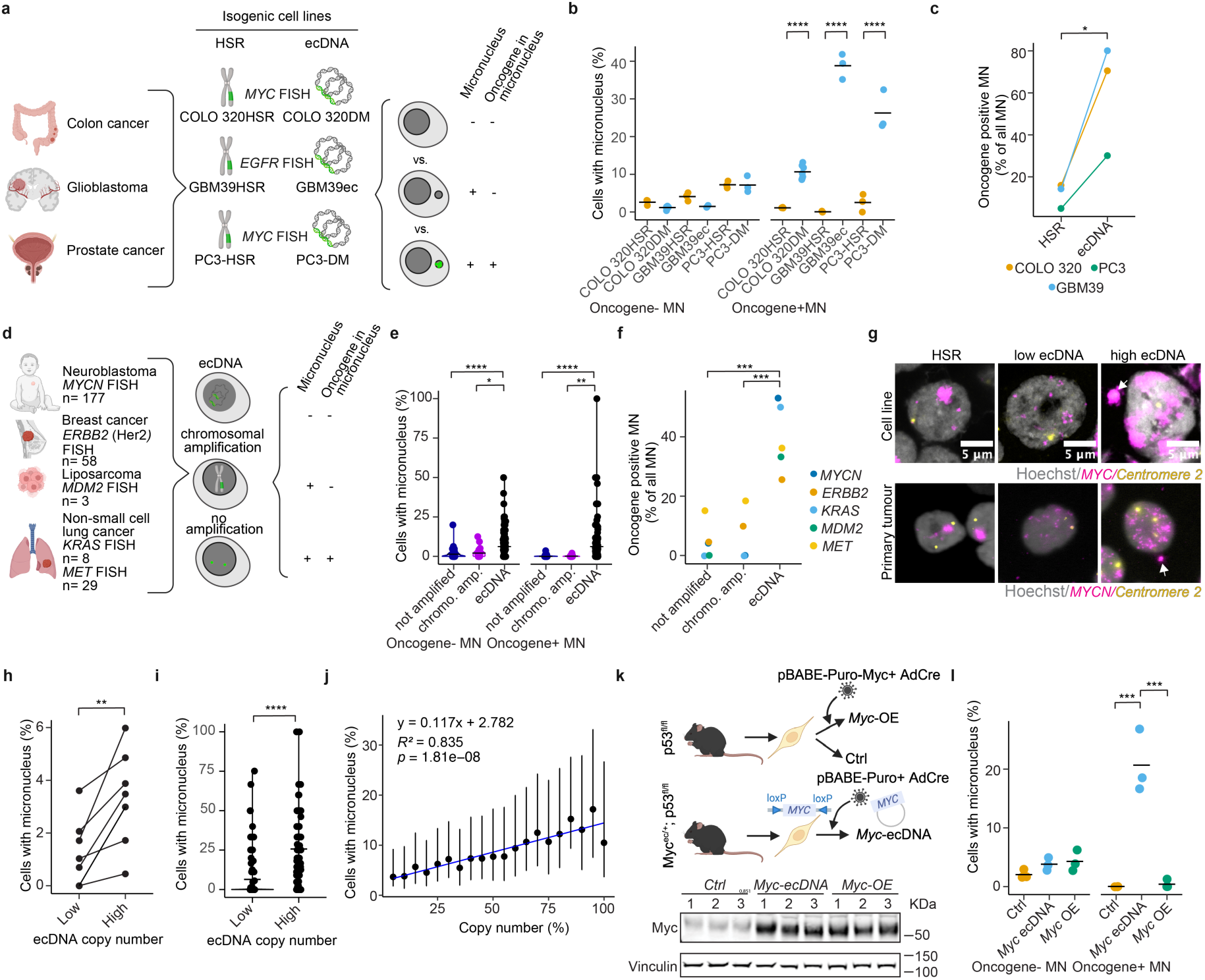
Cancer cells with ecDNA more frequently have micronuclei than cells with chromosomal amplifications. a. Schematic of the experimental design to quantify micronuclei in near-isogenic cancer cell lines. b. Fraction of cells with oncogene-negative and oncogene-positive micronuclei in 3 near-isogenic cell line pairs (n= 6 cell lines), as determined using oncogene-specific FISH. Each dot donates one experimental replicate (n = 2–5 experiments per cell line, n= 37-837 cells per replicate; Student’s *t*-test). c. Fraction of micronuclei containing the amplified oncogene in near-isogenic cell lines (n = 3 isogenic cell line pairs, n= 25-522 micronuclei per cell line). Each dot represents one cell line. d. Schematic of the experimental design to quantify micronuclei in patient biopsies. e. Fraction of cells with oncogene-negative and oncogene-positive micronuclei in primary tumours with ecDNA, chromosomal amplification or without amplification (n = 236 tumours; n= 5-574 cells per tumour sample; Kruskal-Wallis-test), as determined using oncogene-specific FISH. Each dot denotes one patient sample. f. Fraction of micronuclei containing the amplified oncogene across patient cohorts stratified by oncogene (Student’s *t*-test). g. Exemplary micrographs of cell line and *MYCN-*amplified Neuroblastoma tumour sample with an HSR (cell line: COLO 320HSR), low ecDNA content or high ecDNA content (cell line: both COLO 320DM) and micronuclei (arrowheads) stained with Hoechst (grey), *MYCN*-specific FISH (magenta) and chromosome 2 centromeric FISH (yellow). h-i. Fraction of micronucleated cells in the cells with low (bottom 30%, n= 5-192 cells) and high (top 30%, n= 5-192 cells) ecDNA content in cell lines (n = 7 cell lines, COLO 320DM, CHP-212, STA-NB10DM, Tr-14, UKF-NB6, Lan5, SMS-KAN (paired Student’s *t*-test) h) and in primary tumours (n = 148, Kruskal-Wallis-test) (i);). Each dot corresponds to one cell line (h) or one patient sample (i). j. Combined cell line and tumour cell fraction from (h) and (i) with micronuclei and their ecDNA copy number. For each dataset, overall ecDNA copy number as determined by FISH was partitioned into deciles, and micronucleation frequency was calculated within each bin (n = 148 tumours, n= 7 cell lines). k. Experimental design to induce ecDNA in mouse aNSCs (top), and Myc immunoblot (bottom) in three independent *Myc* ecDNA-containing cells (*Myc*-ecDNA) and *Myc*-overexpressing (*Myc*-OE) cells compared to control cells (Ctrl) without ecDNA and not expressing Myc, respectively. l. Fraction of cells with *Myc*-negative (oncogene-negative) and *Myc*-positive (oncogene-positive) micronuclei in *Myc*-ecDNA cells, *Myc*-OE or control aNSCs, as determined by *Myc*-specific FISH and Hoechst staining. Each dot denotes one independent experiment (n = 3; one-way ANOVA).

To rule out high oncogene expression as the cause of increased MN formation, we analysed *p53^fl/fl^;Myc^ec/+^* mouse adult neural stem cells (aNSCs) transduced to express Cre recombinase, which results in engineered *Myc*-containing ecDNAs, compared to *p53fl/fl;Myc^OE^* control cells with chromosomally integrated *Myc* that express comparably high levels of Myc protein (Fig. 1k). Only ecDNA⁺ cells exhibited elevated MN abundance, and these MN were enriched for *Myc* sequences (Fig. 1l). Thus, the presence of ecDNA is linked to elevated MN frequencies in cancer cells.

### Replication stress and DNA damage promote ecDNA cluster detachment and micronucleus formation during mitosis

Because ecDNAs lack centromeres, they can rely on chromosomal tethering to ensure inheritance during mitosis^11,14,27^. We hypothesized that MN formation could occur when ecDNAs fail to tether to chromosomes during cell division (Fig. 2a), as also evidenced by recent reports^14,28^. If ecDNAs detached and mis-segregated stochastically and remained independent, individual ecDNA molecules would be expected to segregate into separate micronuclei and be inherited by both daughter cells. Instead, live-cell imaging of unperturbed COLO 320DM cells expressing fluorescently labelled *MYC* ecDNA (tetR–mNeonGreen) and chromatin (H2B–emiRFP670)^27^ revealed that ecDNA detach from chromatids as clusters during mitosis and are subsequently encapsulated on average into single MN in daughter cells (Fig. 2a-b, Supplementary Video 1). The frequency of detached clusters positively scaled with the abundance of ecDNA in cells (Fig. 2c–d), suggesting that while individual detachment events may occur stochastically, ecDNA coalescence and subsequent mis-segregation are governed by non-random processes.

**Fig 2.**
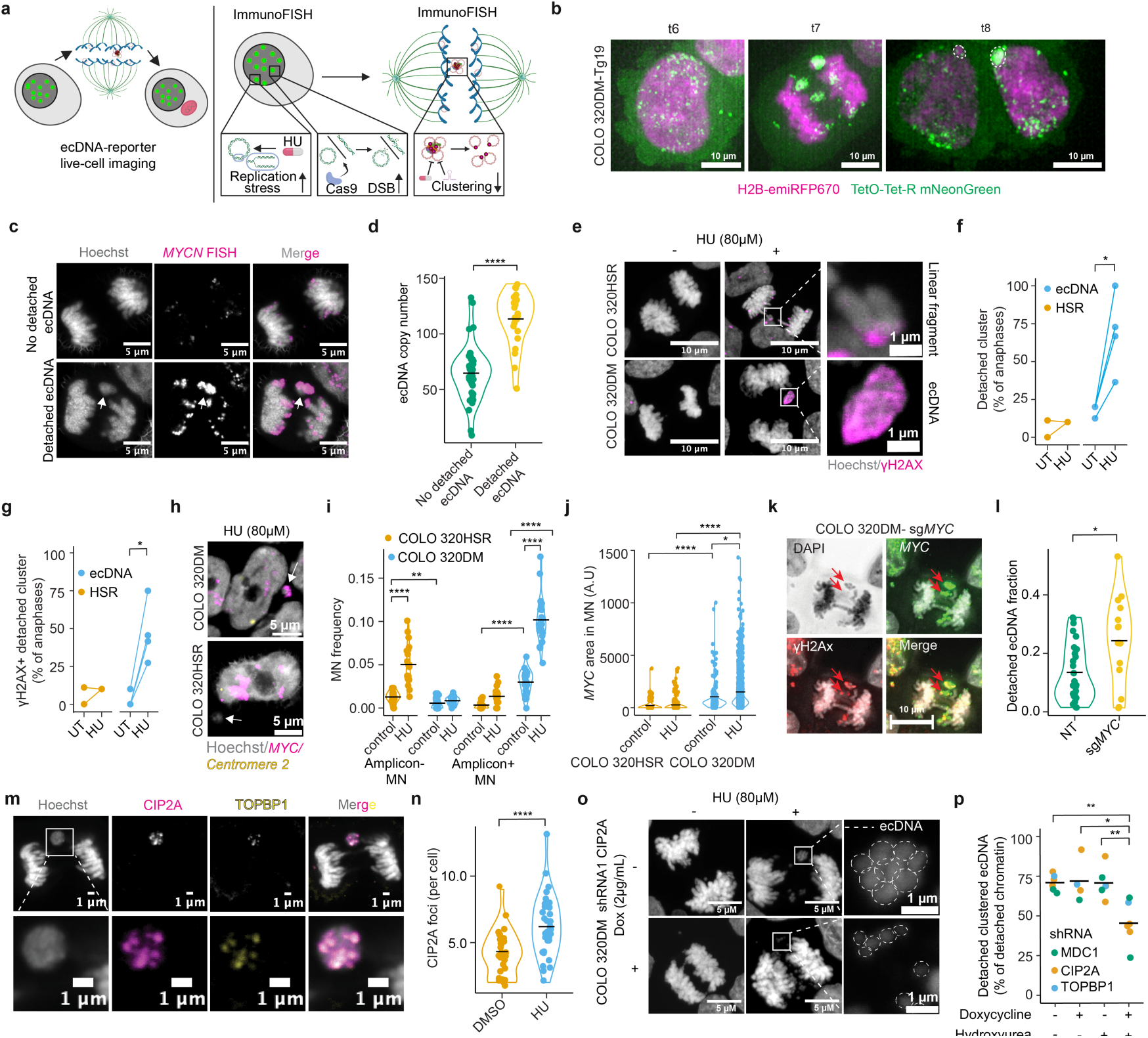
Replication stress induces MDC1-CIP2A-TOPBP1-associated clustered detachment and micronucleation. a. Schematic of experimental design. Live-cell imaging of COLO 320DM-Tg19 cells with fluorescently tagged ecDNA was used to evaluate detachment and micronucleation of ecDNA. FISH combined with immunofluorescence (ImmunoFISH) in fixed interphase and anaphase cells was used to measure replication stress, DNA damage and MDC1-CIP2A-TOPBP1 on ecDNA clusters in mitosis following genetic and pharmacologic perturbations. b. Exemplary photomicrographs taken during live cell imaging of a COLO 320DM-Tg19 cell engineered to visualize *MYC* ecDNA (TetO-TetR-mNeonGreen) during mitotic transition. Cell with ecDNA (green) within its primary nucleus (magenta) before mitosis (left), during anaphase (middle) and after mitosis (right). Dotted lines indicate micronuclei, which were confirmed to be outside the primary nucleus. Scale bar: 10µm. c. Exemplary micrographs of fixed, untreated CHP212 cells that contain *MYCN* ecDNA during anaphase with (bottom) or without (top) detached ecDNA cluster, stained with Hoechst (grey) and *MYCN* DNA FISH (magenta). d. Number of ecDNA copies per cell in CHP212 cells with detached ecDNA clusters during anaphase in untreated cells compared to those cells without such detached clusters. Each dot denotes one anaphase (n = 52; Student’s *t*-test). Black line indicates average in all plots. e. Exemplary photomicrographs of representative fixed COLO 320DM (contains *MYC* as ecDNA, bottom) and COLO 320HSR (contains *MYC* as HSR, top) cells in anaphase that were incubated in the presence or absence of HU (80µM) for 3 days and stained with Hoechst and for γH2AX. Scale bar: 5µm f. Frequency of DNA clusters during anaphase as measured using Hoechst staining of fixed cells in cell lines with ecDNA (n= 4 independent cell lines) compared to cell lines with HSRs (n=2 independent cell lines) in the presence (HU) or absence (UT) of hydroxyurea (80 µM) (n=10-15 anaphases per cell line and treatment group). Each dot represents the average fraction of cells with a DNA cluster per cell line. g. Frequency of detached chromatin clusters positively staining for γH2AX in cell lines containing ecDNA (n=4 independent cell lines) compared to cells with HSRs (n=2 independent cell lines) incubated in the presence or absence of hydroxyurea (80 µM, 24h) (n=10-15 anaphases per cell line and treatment group were assessed). Each dot represents the average per cell line. h. Exemplary photomicrographs of representative fixed COLO 320DM containing *MYC* ecDNA and COLO 320SHR containing *MYC* HSR, stained with *MYC* FISH after 3d HU (80µM). White arrows indicate MN. Scale bar: 5µm i. Fraction of COLO 320DM and COLO 320HSR cells with oncogene-versus oncogene^+^ MN based on oncogene-specific FISH in the presence or absence of HU (50 µM; n= 20-26 MN; Two-way ANOVA). Each dot represents one image. j. Quantification of *MYC* FISH signal in micronuclei in the presence of HU (3d, 50µM) in COLO 320DM compared to COLO 320HSR cells (n = 100–923 MN; Wilcox test). Each dot denotes one MN. k. Exemplary photomicrographs of COLO 320DM cells after CRISPR-Cas9 mediated cutting of *MYC,* which is present on ecDNA in these cells, and stained with DAPI (grey), *MYC* FISH (green), γH2AX immunofluorescence (red) as well as a merged image. l. Fraction of detached ecDNA in anaphase, measured using *MYC* FISH in COLO 320DM cells 24hrs after nucleofection of Cas9 and a guide RNA targeting *MYC (*sg*MYC,* n=15) compared to cells expressing a scrambled sgRNA control (sgNT, n=27) (Student’s t-test). m. Exemplary photomicrographs of a COLO 320DM cell with ecDNA in anaphase (top) as well as an enlarged view of DNA cluster (bottom) stained with Hoechst (grey) and immunofluorescence for TOPBP1 (yellow) or CIP2A (magenta). n. Quantification of immunofluorescent CIP2A foci in COLO 320DM anaphase cells after incubation in the presence of HU (100 μM) for 24hrs compared to untreated cells (n = 31–33 anaphases; Student’s t-test). Each dot denotes one anaphase. o. Exemplary photomicrographs of COLO 320DM cells transduced to inducibly express an shRNA targeting CIP2A in the presence (bottom) or absence (top) of doxycycline (2 μg/ml, 2 days) and in the presence (right) or absence (left) of HU (80 μM, 24 h) in anaphase. Scale bar: 5 μm. Dotted circles represent outlines of small chromatin bodies representing ecDNA. p. Fraction of clustered chromatin as measured using Hoechst staining in cells harbouring ecDNA and inducibly expressing shRNAs targeting MDC1 (two independent sgRNAs, green), TOPBP1 (1 shRNA, blue), or CIP2A (3 independent shRNAs, orange) in the presence or absence of either doxycycline (2µg/ml, 2 days) and/or HU (80µM, 24h) in COLO 320DM cells (n = 10–12 anaphases per condition; one-way ANOVA). Each dot denotes the mean per anaphase. Line indicates the mean.

ecDNAs are frequently damaged and prone to replication stress^29,30^, prompting us to test whether DNA damage promotes clustered detachment and micronucleation. Consistent with these prior findings, significantly higher signs of replication stress and DNA damage were observed on ecDNA compared to HSR (Extended Fig. 3a-f). Treatment with hydroxyurea, which induces replication stress and causes DNA damage by depleting nucleotides, increased signs of replication stress especially on ecDNA (Extended Fig. 3a-f). Nanopore-based fork progression analysis confirmed reduced nucleotide incorporation and disproportional replication fork slowing on ecDNA both in untreated conditions and upon HU (Extended Fig. 4). This resulted in substantially more untethered, γH2AX-positive ecDNA⁺ clusters during anaphase in COLO 320DM cells compared to their isogenic HSR counterparts as well as in non-isogenic cell line pairs (Fig. 2e-g, Extended Fig. 5). As a result, ecDNA⁺ cells exhibited increased MN frequency, and MN were significantly enriched for ecDNA oncogene sequences (oncogene⁺ MN) to an extent exceeding expectations from random segregation (Extended Fig. 6a-b). The increase in MN abundance was rescued by transfection of nucleotides, confirming that HU-induced nucleotide depletion promoted MN formation (Extended Fig. 6c-d). No enrichment of MN for oncogene sequences was observed in cells with chromosomal amplifications, in which the overall MN rate increased following hydroxyurea, but those MN did not contain oncogene sequences (oncogene^-^ MN, Fig. 2h–j, Extended Fig. 6e-f). The frequency of MN after hydroxyurea scaled with the abundance of ecDNA in cells (Extended Fig. 6g-h). As in untreated cells, despite cells harbouring high numbers of the same ecDNA species, each daughter cell typically contained only a single MN (mean 1.03 and 1.19 per cell for CHP212 and COLO 320DM, respectively), suggesting that detached ecDNAs coalesce into a single cluster before segregation and micronucleation in one daughter cell. Targeted CRISPR–Cas9 cleavage of ecDNA reproduced this effect, producing untethered ecDNA clusters that segregated into MN (Fig. 2k–l, Extended Fig. 7), suggesting that damage on ecDNA promotes clustered detachment and micronucleation. Thus, ecDNA is more prone than chromosomal amplifications to respond to replication stress with clustered detachment in mitosis, resulting in micronucleus formation.

The MDC1–CIP2A–TOPBP1 complex stabilizes damaged chromosomal fragments during mitosis^31,32^, raising the possibility that it mediates ecDNA clustering following their damage. Immunofluorescence staining revealed that a subset of detached clusters colocalised with CIP2A, and TOPBP1 (Fig. 2m), with overall signal intensity increasing after hydroxyurea treatment (Fig. 2n, Extended Fig. 8a-b). Pharmacological reduction of CIP2A expression using TD19 dispersed Hoechst-stained clusters into individual foci in ecDNA+ cells, and increased the number of small MN per cell without changing the overall fraction of micronucleated cells (Extended Data Fig. 8c-h). Similarly, shRNA-mediated knockdown of CIP2A, TOPBP1, or MDC1 diminished chromatin clustering during anaphase, while not changing overall detachment frequency (Fig. 2o-p, Extended Fig. 8i-j). Together, these results indicate that the MDC1–CIP2A–TOPBP1 complex is at least in part required for the coalescence of damaged ecDNAs into clusters, similar to what was described for pulverized DNA during chromothripsis^31,32^, which are then inherited as a single MN in daughter cells. This pathway links replication stress to elevated ecDNA mis-segregation and micronucleus formation.

### Single-micronucleus sequencing reveals that multiple distinct ecDNAs tend to coalesce into the same micronucleus

Up to 25% of ecDNA⁺ tumours harbour more than one ecDNA species^27^. Given damaged ecDNAs cluster before the end of mitosis, individual micronuclei might contain oncogenes from distinct ecDNA species. At the same time, DNA damage could in principle generate chromosomal fragments that co-segregate into MN together with attached ecDNA. To determine the genomic content of MN in cancer cell lines with multiple ecDNA species, we developed a laser-microdissection method to isolate single MN and their corresponding primary nuclei, followed by paired-end sequencing (Fig. 3a). This approach was applied to TR-14 cells, which harbor multiple ecDNA species, each carrying a distinct oncogene (eg. *MYCN*, *MDM2*, or *CDK4*) alongside additional passenger genes. The circular structure of these ecDNA was previously assessed by sequence reconstruction^33^, and their extrachromosomal presence, oncogene content and copy number per cell was confirmed by metaphase-FISH (Fig. 3b). After hydroxyurea treatment to induce replication stress, 70% of micronuclei were strongly enriched in all ecDNA species (Fig. 3c; Extended Fig. 9a-b), consistent with collective micronucleation of separate ecDNAs after replication stress. Notably, the majority of ecDNA⁺ MN (92.3%) lacked detectable linear chromatin fragments, arguing against frequent co-segregation of chromosomal fragments together with ecDNA and indicating that ecDNA mis-segregates more readily than chromosomal DNA under replication stress (Fig. 3d-e; Extended Fig. 9c). Importantly, ecDNA was markedly enriched relative to linear chromosomal DNA in micronuclei compared with the corresponding primary nuclei, a pattern that would not be expected if ecDNA mis-segregated stochastically, in which case similar relative enrichments would be anticipated in both compartments. Copy number proportions among ecDNA species within individual MN mirrored those in the primary nucleus (Fig. 3f, Extended Fig. 9d), and enrichment of different ecDNA species correlated positively in MN (Fig. 3g-j), in line with the collective micronucleation of structurally distinct ecDNAs.

**Fig 3.**
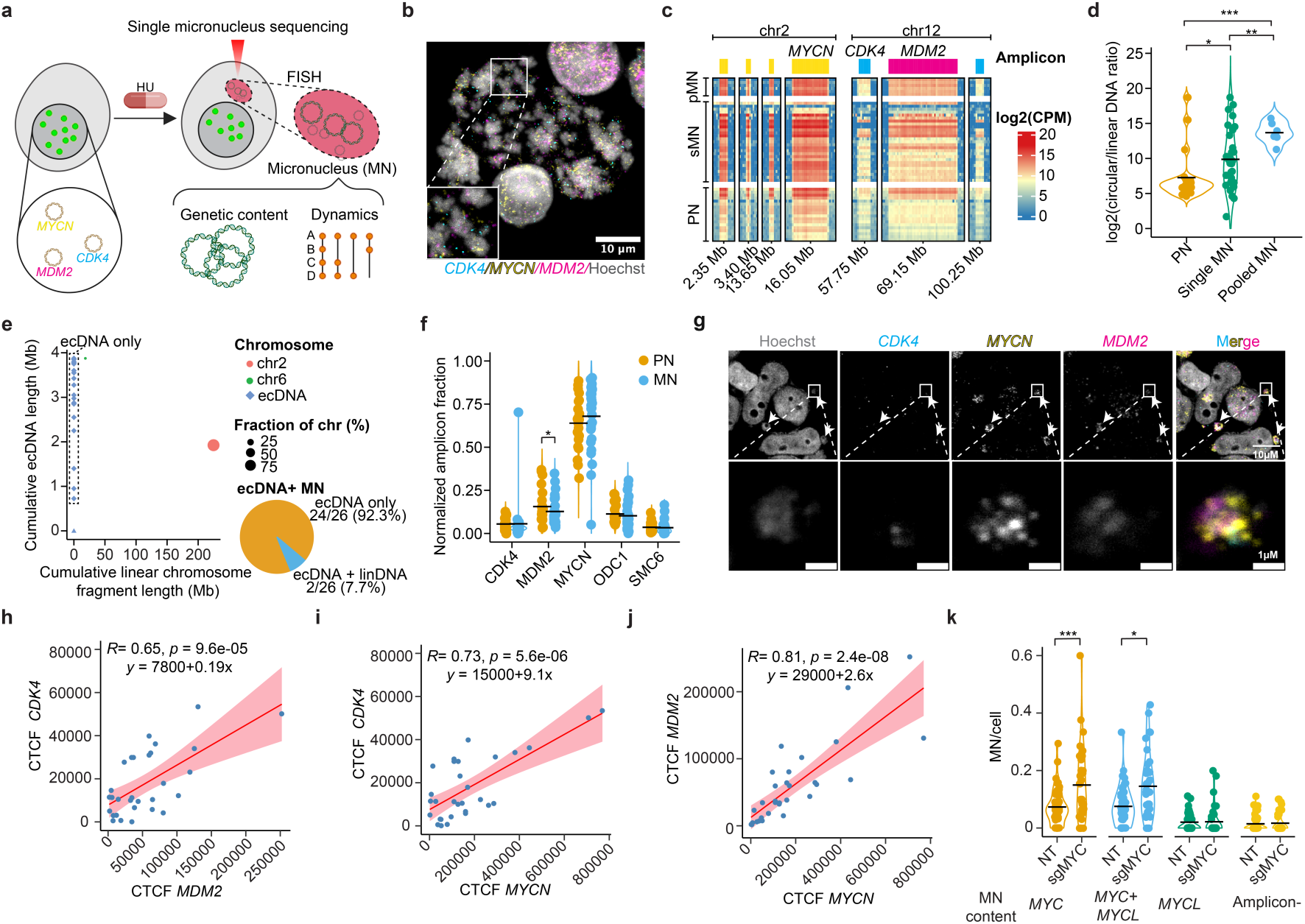
Collective entrapment of distinct ecDNAs in the same micronuclei. a. Experimental design. Laser microdissection (red triangle) was used to isolate micronuclei or primary nuclei, which was followed by sequencing. FISH was used to validate findings. b. Exemplary photomicrograph of a metaphase spread of a TR-14 cell stained with Hoechst (grey) and FISH for oncogenes (*CDK4,* blue; *MYCN,* yellow; *MDM2,* red) present on distinct ecDNA species. c. Laser microdissection directed sequencing of primary nuclei (PN) and micronuclei (MN) after incubation of TR-14 cells in the presence of HU (80 µM). Counts per million (CPM), normalized and log2 transformed read coverage (25kb bins) across ecDNA (*CDK4,* blue; *MYCN,* yellow; *MDM2,* red) and their flanking regions (100 kb; primary nucleus, PN, n = 18; single micronucleus, sMN, n = 27; pooled micronuclei, pMN, n = 6). d. Log2-transformed mean coverage ratio of circular reads over the winsorized mean of all linear reads in PN, sMN and pMN (Welch’s t-test). e. Cumulative ecDNA and linear chromosome length of enriched fragments in single micronuclei. Size of linear chromosomal fragments (dots) denotes fraction of the incorporated whole chromosome or chromosomal fragment. Bottom right: fraction of ecDNA positive MN containing either only ecDNA or ecDNA and linear chromatin. f. Amplicon-size normalized fraction of circular reads mapping to specific ecDNA species in primary nuclei (n=18) and micronuclei (n=27) (Permutation test, 10000 permutations, one-sided). g. Exemplary photomicrograph of a micronucleus in the TR-14 cell line harbouring multiple ecDNA, stained for *CDK4, MYCN* and *MDM2* using FISH. h-j. Correlation between normalised corrected total cell fluorescence (CTCF) from FISH in micronuclei for *CDK4* and *MDM2* (h), *CDK4* and *MYCN* (i), *MDM2* and *MYCN* (j). k. Frequency of micronuclei per cell in DMS273 cells 24h after expression of Cas9 and sgRNA targeting *MYC* (sgMYC) or a scrambled sgRNA (NT) and stained with DAPI and oncogene-specific FISH for *MYC* or *MYCL* to classify MN according to their ecDNA content (n= 38-48, Wilcox test). Each dot represents one image.

To test whether damage to one ecDNA species can result in collective micronucleation of other ecDNA species, we introduced CRISPR–Cas9-mediated double-strand breaks on *MYC* ecDNA of DMS273 cells, which also harbour a separate ecDNA species encoding for *MYCL* (Extended Fig. 10a). Targeted cleavage increased γH2AX foci on *MYC* and promoted its presence in MN, while concurrently enriching *MYCL* ecDNA in the same MN (Fig. 3k, Extended Fig. 10b-f).

Taken together, these findings suggest that ecDNA mis-segregate into micronuclei *en masse* in cells with elevated ecDNA damage and is largely independent of linear chromatin fragments, raising the possibility that this impacts ecDNA inheritance patterns (Extended Fig. 11).

### Micronucleation leads to asymmetric inheritance of ecDNA with reduced oncogenic transcription

Live-cell imaging revealed that ecDNA clusters, both under basal conditions and following hydroxyurea treatment, segregated on average into a single micronucleus (MN) and one of the two daughter cells, suggesting that micronucleation introduces asymmetry into ecDNA inheritance (Extended Fig. 12a-b, Supplementary Video 2-3). This suggests that ecDNA micronucleation skews previously observed random inheritance patterns^11^, which would have implications for tumour evolution. To quantify this effect, we measured ecDNA copy number in paired daughter cells at cytokinesis using DNA FISH after hydroxyurea treatment. Compared to untreated controls, hydroxyurea markedly increased the degree of ecDNA imbalance between daughter cells (Extended Fig. 12b), confirming that replication stress promotes asymmetric segregation through micronucleation.

To further investigate the consequence of asymmetric segregation for the inheritance of ecDNA, we developed a computational model (Extended Fig. 12c). Assuming a probability *q* of asymmetric segregation and probability 1-*q* of random segregation. The resulting ecDNA distribution in daughter cells shifted from unimodal (random segregation) to bimodal or trimodal patterns as *q* and 𝑝_1_ varied (Extended Data Fig. 12d). Using an approximate Bayesian computation framework (ABC), we inferred optimal parameter estimates for *p*_1_ and *q* (Extended Data Fig. 11e-i). Applying this to our experimental data from pairs of daughter cells with and without hydroxyurea treatment, we observed increased biased segregation post-treatment (Extended Fig. 12j-l). Thus, replication stress enhances ecDNA segregation asymmetry (Fig. 11).

Linear chromatin fragments incorporated in MN remain epigenetically and transcriptionally silenced^21–23^. We set out to test if ecDNA is also repressed in MN following replication stress (Extended Fig. 13a). During mitosis, ecDNA retained high H3K27Ac and low H3K27me3, much unlike chromosomal DNA, which showed the expected inactive chromatin patterns (Extended Fig. 13b-c). However, ecDNA in MN had significantly reduced H3K27Ac and stably low H3K27me3 compared to ecDNA in mitosis and the primary nuclei following hydroxyurea treatment (Extended Fig. 13d-e). Chromatin immunoprecipitation sequencing confirmed reduced H3K27ac at ecDNA loci, including *MYC* (Extended Fig. 13f). Micronuclei also exhibited significantly lower levels of pRNAPII-S5, indicating reduced transcriptional activity in micronuclei (Extended Fig. 13g-h). Indeed, intron RNA-FISH further confirmed that most micronuclei in ecDNA-containing cell lines lacked detectable oncogene expression, independent of the oncogene encoded (Extended Fig. 13i). RNA sequencing confirmed the decreased expression of ecDNA-encoded genes, including *MYC*, alongside reduced MYC-driven transcriptional programs (Extended Fig. 13j-l). Collectively, these results show that, unlike its high expression in the nucleus and throughout mitosis^27,28^, ecDNA within micronuclei is transcriptionally silenced, suggesting that micronucleation restrains the oncogenic potential of ecDNA.

### ecDNA micronucleation impacts cancer cell fitness and is associated with improved patient survival

To investigate the functional consequences of ecDNA⁺ micronuclei, we performed live-cell imaging under both untreated conditions and following hydroxyurea exposure to track daughter cells inheriting a micronucleus after mitosis (Fig. 4a). Cells that inherited large, ecDNA-containing MN (> 2 µm diameter) showed delayed progression to the next division, and higher death rates than the daughter cell lacking a MN (Fig. 4b–c; Extended Fig. 14a-b, Supplementary video 4-6). These effects were independent of ecDNA copy number in the primary nucleus (Extended Fig. 14c-d), but were strongest when MN contained high ecDNA content, indicating that ecDNA in MN impose a direct fitness penalty.

**Fig. 4:**
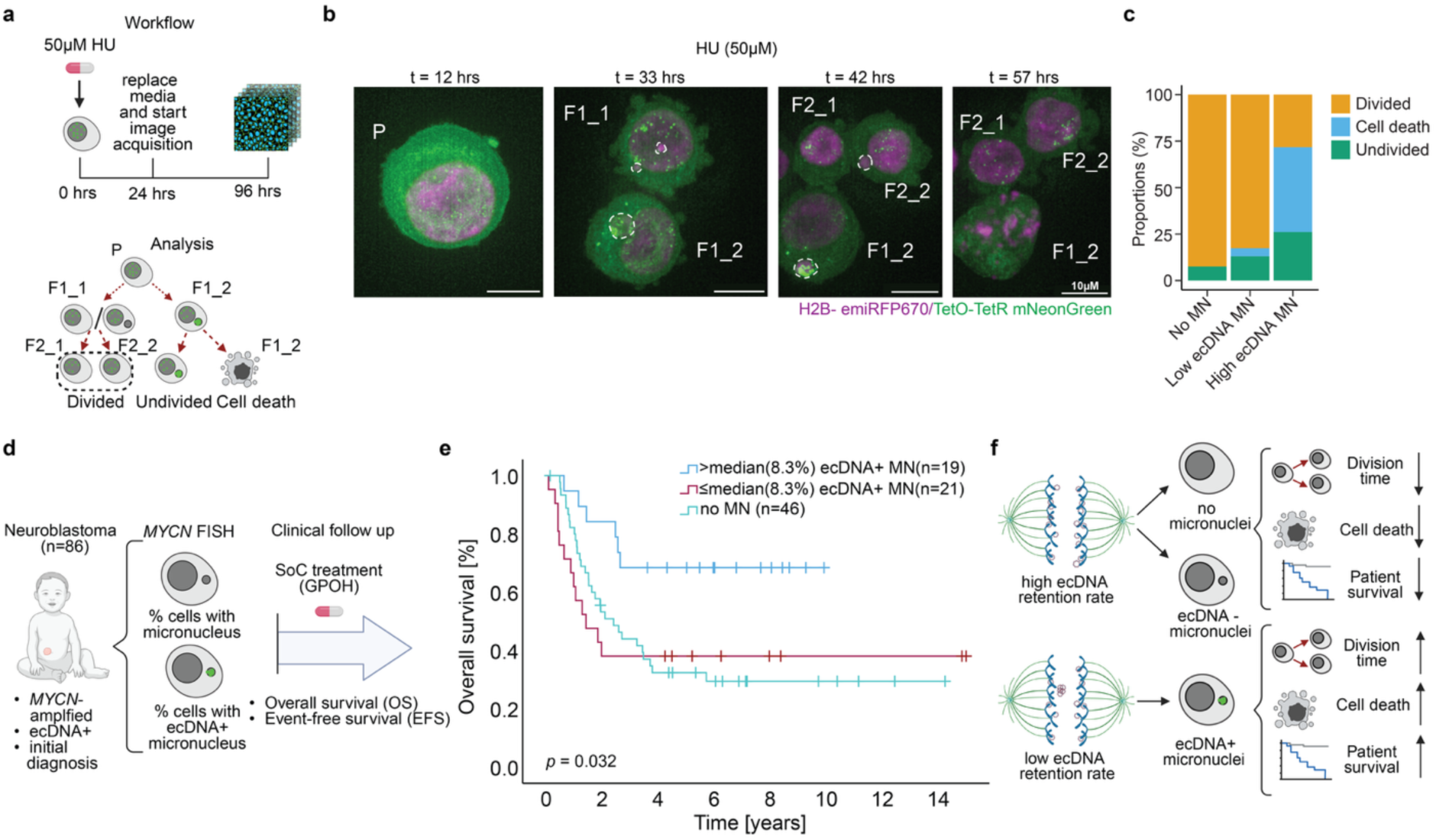
The presence of ecDNA in micronuclei is associated with reduced cancer cell fitness and improved patient outcome. a. Experimental design for live-cell imaging and analysis. Cells were tracked for two generations, from the parental cell (P), to generation F1, daughter cells (F1_1 and F1_2) to generation F2, granddaughter cells (F2_1 and F2_2). b. Representative frames from time-lapse imaging of a COLO 320DM-Tg19 parental cell (P, left) undergoing cell division twice (F1 and F2 generations) in the presence of HU (50µM), in which *MYC* ecDNA (green) and chromatin (magenta) are fluorescently labelled. Micronuclei are marked with dotted lines and were confirmed in 3D reconstructions to be outside the nucleus. Cell with ecDNA-containing MN (F1_2) exhibits nuclear blebbing at t=57h, indicative of cell death. Scale bar: 10 μm. c. Quantification of COLO 320DM-Tg19 cell fates as determined using live cell imaging in the presence of HU (50µM) for cells without micronuclei (No MN), with MN containing low ecDNA content (below median; low ecDNA MN), or with MN containing high ecDNA content (above median; high ecDNA MN). Cell fates include cell division, cell death, or remaining undivided (n = 143 individual cells at the end of the 72h imaging period). d. Schematic overview of the analysis of 86 *MYCN* ecDNA-positive neuroblastoma biopsies, in which the presence of micronuclei and their content was determined using *MYCN* FISH and DAPI staining (registered in the trials NB97, n = 2; NB04, n = 67; NB2016, n = 17). Median follow-up, 6.8 years (range 0.14–15.9) for 36 survivors. e. Kaplan–Meier survival curves for patients with more than 8.3% cells (median) in their biopsy with ecDNA-containing micronuclei (ecDNA⁺ MN), patients with less than 8.3% ecDNA⁺ MN, or those patients without MN (no MN; log-rank test). f. Summary schematic of the main findings.

If this penalty constrains tumour growth, patients whose tumours contain abundant ecDNA⁺ MN should have less aggressive disease. We examined diagnostic biopsies from 387 patients and selected 86 children with *MYCN*-amplified neuroblastoma in which ecDNA was identified by FISH before treatment (Fig. 4d, Extended Fig. 15a). For each case, we quantified the proportion and ecDNA content of MN at baseline and compared these with clinical outcome after standard-of-care treatment. Neither total ecDNA copy number in the primary nucleus nor overall MN frequency correlated with survival (Extended Fig. 15b-g). In contrast, tumours with a high fraction of ecDNA⁺ MN exhibited significantly longer event-free and overall survival than tumours with few or no ecDNA⁺ MN (Fig. 4e; Extended Data Fig. 15h). Modelled as a continuous variable and assuming linearity, ecDNA⁺ MN frequency remained independently prognostic (Overall survival: hazard ratio 0.951, 95% CI 0.914–0.990, *p* = 0.015, event-free survival: hazard ratio 0.956, 95% CI 0.923–0.990, *p* = 0.012), corresponding to a 4.9 % reduction in risk of death for every 1 % increase in ecDNA⁺ MN. These findings indicate that an increase in micronuclei containing ecDNA (ecDNA⁺ MN) is associated with impaired proliferative capacity of cancer cells and reduced tumour fitness (Fig. 4f).

Our results support the view that faithful inheritance of ecDNA depends on chromosomal tethering during mitosis^13,14,34^. When this process fails, oncogene-bearing ecDNAs can coalesce into MDC1–CIP2A–TOPBP1–stabilized clusters that segregate asymmetrically and are sequestered into micronuclei, a process that is potentiated by replication stress–induced damage. This asymmetric accumulation of ecDNA within micronuclei is associated with decreased oncogene output, impaired daughter-cell fitness and corresponds to improved survival in children with *MYCN*-amplified neuroblastoma.

Together, these findings link ecDNA^+^ micronucleus formation to reduced tumour fitness, including at the patient level. The observation that sequestration of oncogene-containing ecDNA into micronuclei is associated with a proliferative penalty and improved event-free and overall survival in children with neuroblastoma highlights the functional relevance of ecDNA-containing micronuclei and suggests their potential utility as biomarkers for patient stratification and for therapeutic strategies aimed at selectively damaging ecDNA.

## Methods

### Cell lines

Human tumour cell lines were obtained from the American Type Culture Collection (ATCC) or a gift from collaborative laboratories. The identity of all cell lines was verified by short tandem repeat genotyping (Eurofins Genomics). The absence of *Mycoplasma* sp. contamination was determined using a Lonza MycoAlert system (Lonza). For this study, 11 ecDNA amplified cell lines (DMS273, STA-NB10DM, COLO320 DM, TR-14, CHP-212, Lan5, UKF-NB6, KP-N-YN, PC3-DM, SiMA, SMS-KAN), 5 HSR amplified cell lines (STA-NB10HSR, COLO 320HSR, PC3-HSR, NGP, IMR5/75), 2 non-amplified cancer cell line (ShEP, HeLa) and 1 non-cancerous cell line (RPE-hTERT) were used. HEK293T cells were used for virus production. Near isogenic cell line pairs GBM39ec, GBM39HSR, COLO320 DM, COLO320 HSR, PC3-DM, PC3-HSR cells were extensively characterized before^29^.

### Cell culture

Most neuroblastoma and colon cancer cell lines were cultured in RPMI-1640 medium (Gibco) supplemented with 1 % of penicillin, streptomycin, and 10 % of fetal calf serum (FCS) (Thermo Fisher). STA-NB10DM and HSR cells were cultured in RPMI-1640 medium (Gibco) supplemented with 1 % of penicillin, streptomycin, 10 % of fetal calf serum (FCS) (Thermo Fisher), 1 % Glutamax (Thermo Fisher), 1 % Sodium Pyruvat (Thermo Fisher) and 10 mM HEPES (Sigma). UKF-NB6 was cultured in IMDM medium (Gibco) supplemented with 1 % of penicillin, streptomycin, and 10 % of FCS. RPE and HEK293T cells were cultured in DMEM (Gibco) supplemented with 1 % of penicillin, streptomycin, and 10 % of FCS. GBM39ec and GBM39HSR cell lines were cultured in Dulbecco’s Modified Eagle’s Medium/F12 (Gibco, catalogue no. 11320-033) supplemented with 1× B27 (Gibco, catalogue no. 17504-01), 20 ng/ml epidermal growth factor (Sigma, catalogue no. E9644), 20 ng/ml fibroblast growth factor (Peprotech, catalogue no. AF-100-18B), 5 µg/ml heparin (Sigma, catalogue no. H3149) and 1× GlutaMAX (Gibco, catalogue no. 35050-061). COLO 320DM, COLO 320HSR, PC3-DM, PC3-HSR and COLO 320DM-Tg19 (live-cell imaging line) cells were cultured in DMEM (Corning, #10-013-CV) with 1% PSQ and 10% FBS. To assess the number of viable cells, cells were trypsinized (Thermo Fisher), resuspended in medium, and sedimented at 300 *g* for 5 minutes. STA-NB10 cells were split using Accutase (Thermo Fisher). HeLa cells were cultured in Advanced DMEM-F12 (Gibco) with addition of 10% FCS, 1% Penicillin-Streptomycin (Gibco) and 1% GlutaMax (Gibco). Cells were then resuspended in medium, mixed in a 1:1 ratio with 0.02 % trypan blue (Thermo Fisher), and counted with a Bio-Rad TC20 cell counter. CHP212-PIP-FUCCI (Nanopore-based replication fork dynamics) cells were generated by lentiviral transduction of pLenti-PGK-Hygro-PIP-FUCCI (Addgene # 138714). Transduced cells were selected by FACS sorting 10 days after transduction and kept under 100 μg/mL Hygromycin B selection.

In drug experiments, cells were treated for 24 or 72h with 50µM or 80 μM or 100µM hydroxyurea or a combination of 80 μM hydroxyurea together with 1 μM TD-19 (Merck Millipore, 532912) as indicated in figure legends respectively. Control cells were treated with an equivalent volume of DMSO or H2O. Drug concentrations were selected based on cell viability as assessed using the CellTiter-Glo Assay (Promega G7571). GBM39ec/GBM39HSR cells were treated with 100 μM hydroxyurea for 5 and 72 hrs, EdU (10 μM) was added for 30 mins before fixing. dNTP supplementations were conducted at the indicated concentrations at a 1:1 ratio with Lipofectamine 3000 (ThermoFisher, L3000001) according to manufactures protocol.

### Genetically induced ecDNA cell lines

Adult Neuronal Stem Cells (aNSCs) were isolated from Mycec/+; p53fl/fl and p53fl/fl animals described in Pradella et al.^35^ and cultured following the protocol described by Ahmed et al.^36^ aNSC were maintained in laminin-coated (Sigma-Aldrich, L2020) dishes in NeuroCult Stem Cell Basal Media with NeuroCult Proliferation Supplement (Mouse & Rat) (Stem Cells Technologies, 05702), 20 ng ml−1 EGF (Stem Cells Technologies, 78006), 10 ng ml−1 bFGF (Stem Cells Technologies, 78003), and 2 μg ml−1 heparin (Stem Cell Technologies, 07980). Cells were tested for mycoplasma contamination and maintained in a humidified, 5% CO2 atmosphere at 37 °C. aNSC overexpressing Myc transgene were generated by infecting p53fl/fl aNSC with pBABE-Puro-Myc retrovirus. Mycec/+; p53fl/fl and p53fl/fl aNSC were also infected with pBABE-Puro retrovirus. Upon infection, cells were selected with puromycin (1 μg ml−1; Sigma-Aldrich, P9620) for 3-4 days. Selected cells were then infected with AdCre (ViraQuest, VQ-Ad-CMV-Cre; 1 × 1012particles per milliliter; 091317) to promote p53 loss and Myc-containing ecDNA accumulation as described in Pradella et al.^35^

### Retrovirus production

pBABE-Puro-C-Myc plasmid was generated by site-directed mutagenesis from pBABE-Puro-C-Myc T58A (Addgene #53178) by removing the T to A mutation in the mouse coding region. For retrovirus production, HEK-293T cells were seeded in high-glucose Dulbecco’s modified Eagle medium (DMEM, 4.5 g l−1; Thermo Fisher Scientific, 11995065) supplemented with 10% heat-inactivated fetal bovine serum (FBS, Sigma-Aldrich, F2442) without antibiotics in 100 mm petri dishes. The next day, cells at approximately 60–70% confluence were transfected with 20 µg of retroviral vector carrying the mouse C-Myc cDNA or a control vector, 5 µg of packaging plasmid and 1 µg of envelope plasmid. After 24 h, the medium was replaced with DMEM with antibiotics. Cells were incubated for 48 h with two consecutive collections of the medium containing the retroviral particles at 24 and 48 h. Medium collected was filtered using a 0.45 µm filter unit and used to transduce aNSC cells by incubating cells with viral supernatant supplemented with polybrene (0.2 µl ml−1; Sigma, TR-1003G).

### Antibodies for immunoblots

For immunoblots of aNSCs the following primary antibodies were used: anti-vinculin (1:5000; Millipore, MAB3574), anti-c-MYC (1:000; Cell Signaling, D84C12). The following secondary antibodies were used: IRDye 800 anti-rabbit (LICOR, 926-32213) and IRDye 680 anti-mouse (LICOR, 926-68072). For shRNA knockdown conformation, the following antibodies were used: Anti-CIP2A (1:5000, Santa Cruz, sc-80659), anti-TOPBP1 (1:5000, Sigma-Aldrich, ABE1463), anti-MDC1(1:5000, abcam, ab11169), anti-vinculin (1:5000, Cell signalling, 4650S), goat anti-Mouse IgG (H+L) Cross-Adsorbed Secondary Antibody, Alexa Fluor™ 647 (1:5000, Invitrogen, A21235) and goat anti-rabbit IgG (H+L), horseradish peroxidase conjugates (1:5000, Invitrogen, G21234) were used.

### Antibodies or immunofluorescent staining

For staining for single-micronucleus sequencing, samples were incubated in mouse anti-pan-histone H11-4 (1:500; Merck, MAB3422), followed by incubation with Alexa555 (1:1000; Invitrogen, A-31570) or Alexa488 anti-mouse secondary antibody (1:1000; Invitrogen, A-11001). For immunofluorescent staining followed by imaging, the following primary antibodies were used: anti-pRPA2-S33 (1:1000, Novus Biological, NB100-544), anti-Lamin B1 Monoclonal antibody (proteintech, 66095-1-Ig, IF - 1:500), Anti-H2AX (1:500/ 1:250, Sigma Aldrich, 05-636), anti-CIP2A (1:500, Santa Cruz, sc-80659), anti-TOPBP1 (1:500, Sigma-Aldrich, ABE1463), anti-pRNAPolII(Serine5, 1:500, Merck, MABE954), anti-H3K27ac (1:500, Active Motif, 39133) or anti-H3K27me3 (1:500, Thermo Fisher, G.299.10). Followed by incubation with secondary antibodies: Alexa Fluor 647, Goat anti-Rabbit IgG (H+L) Cross-Adsorbed Secondary Antibody (1:1000, Invitrogen, A21244) and goat anti-Mouse IgG (H+L) Cross-Adsorbed Secondary Antibody, Alexa Fluor™ 647 (1:1000, Invitrogen, A21235). For ChIP-sequencing, the following antibodies were used: anti-H3K27Ac (1:600, Diagonode, C15410196) and anti-H3K27me3 (1:600, Diagonode, C15410195).

### Immunoblots of aNSC

aNSC cells were lysed in Laemmli buffer, supplemented with protease (cOmplete; Roche, COEDTAF-RO) and phosphatase (EDTA-free Protease Inhibitor Cocktail; Roche, PHOSS-RO) inhibitors. Proteins were separated by SDS polyacrylamide gel electrophoresis and analyzed by western blotting using standard procedures. After protein transfer, the nitrocellulose membranes (Bio-Rad, catalogue no. 1704271) were blocked by incubation with LICOR Intercept (TBS) blocking buffer. Images were acquired using an Odyssey Imaging System (LICOR).

### Imaging of aNSCs

aNSC were fixed with ice-cold methanol for 20 min. Briefly, fixed slides were pretreated with Pepsin (Sigma P-6887), dehydrated and counterstained with DAPI (Thermo, D1306) and mounted with Prolong Diamond Antifade Mounting medium (Thermo, P36965). Images were acquired with a REVOLVE R4 (Echo Laboratories) microscope through a ×63 objective lens. Images were analyzed in the built-in Echo app. Each nucleus was evaluated for the presence of micronuclei and Myc amplification using DNA FISH.

### Western immunoblotting

Whole-cell protein lysates were prepared by lysing cells in 10x Cell lysis buffer (Cell signaling) supplemented with cOmplete (Roche) and PhosStop (Roche) phosphatase inhibitors. Protein concentrations were assessed by bicinchoninic acid assay (BCA, Santa Cruz Biotechnology). For 5 minutes, 15 μg of protein was denatured in Laemmli buffer at 90 °C. Samples were run on NuPage 10 % polyacrylamide, 1 mm Tris-Glycine Protein Gels (Thermo Fisher Scientific) and transferred to PVDF membranes (Roche). Membranes were blocked with 5 % dry milk in TBS with 0.1 % (v/v) Tween-20 (Carl Roth). Membranes were probed with primary antibodies overnight at 4 °C and then with secondary antibodies conjugated to horseradish peroxidase for 1 hour at room temperature. Chemiluminescent detection of proteins was carried out using Pierce ECL Western blotting substrate (Thermo Fisher) and the ChemiDoc (Bio-Rad). Densitometry was performed using ImageJ (NIH).

### Inducible shRNA knock-down plasmid cloning

shRNA constructs were designed using the genetic perturbation platform by the Broad Institute and ordered from Eurofins. They were cloned into pRSITEP-U6Tet-sh-EF1-TetRep-2A-Puro (Celltecta) following the manufacturer’s instructions. We generated three independent shRNAs targeting CIP2A (1: TGCGGCACTTGGAGGTAATTT, 2: ATTTGTGACTTCGTAACAATA, 3: GCTAGTAGACAGAGAACATAA), two targeting TOPBP1 (1: GATAGATTTGGGTAGTAATTT, 2: GCTGCAAAGAAGTGGAATTTA), and one targeting MDC1 (1: CGGACCAAACTTAACCAAGAA). Successful insertion was confirmed by full plasmid sequencing (Plasmidsaurus).

### Lentivirus production and cell transduction

Lentivirus production was carried out as previously described^37^. In brief, HEK293T cells were transfected with PEI MAX (Polysciences Inc) in a 2:1:1 ratio of the lentiviral vector and psPAX2 and pMD2.G packaging plasmids (Addgene), according to the manufacturer’s instructions. Viral supernatant was collected 48 and 72 hours after transfection. The supernatant was pooled, filtered, and stored at –80°C. Cells were transduced with virus particles in the presence of 5 μg/ml polybrene (Merck). Cells were transduced for 1 day in antibiotic-free medium and then grown in full medium for 1 day. Neuroblastoma cells were then selected for 2 days with puromycin hydrochloride (10 μg/ml) or geneticin disulphate (G418, Roth) (2mg/ml). Plasmid expression was induced using 5 μg/ml doxycycline (Sigma-Aldrich Inc., D9891) for 2 days. Western blot validation was done as described above.

### Targeted DNA damage induction by CRISPR-cas9

To induce targeted DNA damage on amplicons, which we refer to as a DNA segment that has undergone focal, high-level amplification, in COLO 320DM/HSR, DMS273 cells, we used CRISPR-cas9. CrRNA used in COLO 320DM/HSR was designed to target an intergenic region amplified in both cell lines (sequence: chr8:+127572615: TACAACGAACTATTTAACCG, TGG(pam)). crRNA used in targeting *MYC* containing ecDNA in DMS273 cells was: GGAGAGCTTGTGGACCGAGC. Non-targeting control was also included (IDT, cat: 1072544). Equal amounts of crRNA and tracerRNA were mixed and annealed before mixing with Alt-R™ S.p. Cas9 Nuclease V3, (IDT, cat:1081059) to make the RNP complex. RNP complex was stabilized by incubating at room temperature for 15 minutes, after which it was delivered into cells using electroporation with the Nucleofector system from Lonza, using program CM-138 and solution SF (Lonza, Catalog #: V4XC-2012). After electroporation, cells were immediately seeded onto coverslips previously coated with fibronectin (10 µg/ml). Cells were fixed at the indicated time points and subjected to multiplex immunofluorescence and DNA-FISH staining.

### Preparation of metaphase spreads

Cells were grown to 80% confluency in a 15 cm dish and metaphase-arrested by adding KaryoMAX™ Colcemid™ (10 µL/mL, Gibco) for 1-2 hours. Cells were washed with PBS, trypsinized and centrifuged at 200 g for 10 min. We added 10 mL of 0.075 M KCl preheated at 37 °C, one mL at a time and vortexing at maximum speed in between. Afterwards, cells were incubated for 20 min at 37 °C. 5 mL of ice-cold 3:1 MeOH/acetic acid (kept at -20 °C) were added, one mL at a time followed by resuspension of the cells by flicking the tube. The sample was centrifuged at 200 g for 5 min. Addition of the fixative followed by centrifugation was repeated four times. Two drops of cells within 200 µL of MeOH/acetic acid were dropped onto prewarmed slides from a height of 15cm. Slides were incubated overnight.

### Fluorescence in situ hybridisation (FISH)

Metaphase spreads or cells seeded on coverslips for interphase FISH were fixed in MeOH/acetic acid for 20 min at -20 °C followed by a PBS wash for 5 min at RT. The wells were removed, and slides were digested in Pepsin solution (0.001 N HCl) with the addition of 10 µl pepsin (1 gr/50 mL) at 37°C for 10 min. After a wash in 0.5x SSC for 5 min, slides were dehydrated by washing in 70 %, 90 % and 100 % cold ethanol stored at -20 °C (3 min in each solution). Dried slides were stained with either a 5 µl of Vysis LSI N-MYC SpectrumGreen/CEP 2 SpectrumOrange Probes (Abbott), ZytoLight ® Spec CDK4/CEN12 Dual Color Probe (ZytoVision) or ZytoLight ® SPEC MDM2/CEN 12 Dual Color Probe (Zytovision), covered with a coverslip and sealed with rubber cement. Denaturing occurred in a Thermobrite (Abbott) for 5min at 72 °C followed by 37 °C overnight. The slides were washed for 5 min at RT within 2× SSC/0.1 % IGEPAL, followed by 3 min at 60 in 0.4× SSC/0.3 % IGEPAL (Sigma-Aldrich Inc.) and further 3 min in 2× SSC/0.1 % IGEPAL at RT. Dried slides were stained with 12 µl Hoechst 33342 (10 µM, Thermo Fisher) for 10 min and washed with PBS for 5 min. After drying, a coverslip was mounted on the slide and sealed with nail polish. Images were taken using a Leica Stellaris Confocal microscope. For ecDNA copy number estimation, we counted foci using FIJI’s find maxima function. Nuclear boundaries were marked as regions of interest. The threshold for signal detection within the regions of interest was determined manually and used for all images analyzed within one group.

### Indirect Immunofluorescence (IF)

Cells were fixed in 4% paraformaldehyde (Electron Microscopy Science) prepared in PHEM buffer (Sigma-Aldrich) for 10 minutes at room temperature, with fresh fixative applied after the first 2 minutes. Fixed cells were washed three times in 125 mM glycine/PHEM for 5 minutes each, followed by permeabilization with 0.5% Triton X-100 (Sigma-Aldrich) in PHEM for 10 minutes at room temperature. After a single 5-minute wash in PHEM, cells were blocked in 5% fetal calf serum (FCS, Sigma-Aldrich) containing 0.1% Tween-20 (Bio-Rad) in PHEM for 30 minutes at room temperature. Primary antibodies were diluted in 5% FCS/0.1% Tween-20/PHEM and applied to cells overnight at 4 °C. The following day, cells were washed three times in 0.1% Tween-20/PHEM for 10 minutes each and incubated with secondary antibodies diluted in 5% FCS/0.1% Tween-20/PHEM for 2 hours at room temperature. After incubation, cells were washed three additional times in 0.1% Tween-20/PHEM for 10 minutes each. Nuclei were counterstained with 5 µM Hoechst 33342 (Thermo Fisher Scientific) in PHEM for 10 minutes, followed by three washes in PHEM for 5 minutes each. Cells were air-dried for 5 minutes and mounted onto SuperFrost glass slides (Epredia) using 5 µl of ProLong Glass Antifade Mountant (Thermo Fisher Scientific). For double immunofluorescence, both primary and secondary antibodies were applied simultaneously.

### Combined FISH and immunofluorescence (ImmunoFISH)

Interphase cells were plated on Poly-D-lysine-coated coverslips (Neuvitro Corporation) in 24-well plates. Cells were fixed with 4% paraformaldehyde for 10 minutes at room temperature, followed by three washes with 125 mM glycine in PBS (5 minutes each). Permeabilization was performed using 0.5% Triton X-100 in PBS for 10 minutes at room temperature, followed by three washes with PBS. Cells were then blocked in 3% bovine serum albumin (BSA) for 30 minutes and incubated overnight at 4 °C with the primary antibody diluted in 3% BSA. After three additional PBS washes, cells were incubated with the secondary antibody (diluted in 3% BSA) for 2 hours at room temperature. Post-incubation, cells were re-fixed in 4% paraformaldehyde for 20 minutes at room temperature. DNA was subsequently denatured by treating cells with 0.7% Triton X-100 and 0.1 M HCl on ice, followed by additional chemical denaturation in 1.9 M HCl for 30 minutes at room temperature. The slides were then washed with 125 mM glycine in PBS, followed by a 5-minute wash in 2× saline-sodium citrate (SSC) buffer. Dehydration and probe hybridization was conducted following the standard FISH protocol described above.

### Triple FISH

Cells were cultured on poly-D-lysine–coated coverslips (Neuvitro Corporation) placed in 24-well plates and fixed in methanol/acetic acid (3:1, v/v) for 20 min at −20 °C. Coverslips were equilibrated in 2× SSC and subsequently dehydrated through a graded ethanol series (70%, 85%, and 100%, 2 min each). For hybridization, 2 µl of each probe specific for *CDK4, MYCN*, and *MDM2* (myTags®, Arbor Biosciences) conjugated to Alexa 488, Atto 550, and Atto 633, respectively, were mixed with 10 µl hybridization buffer (50% formamide, 10% dextran sulfate, 40 ng/µl RNase A in SSCT). A 5 µl aliquot of this mixture was applied per coverslip. Hybridization was carried out at 80 °C for 10 min followed by incubation at 37 °C overnight. Post-hybridization washes were performed according to the standard protocol for single-gene DNA FISH. For ecDNA copy number estimation in triple FISH experiments, foci were detected using Radial Symmetry-FISH (RS-FISH)^38^. Parameters were set manually for each channel and applied to all images from the same coverslip.

### Live-cell imaging

Live-cell imaging presented in Extended Fig. 10a was performed as previously described^27^. The stably engineered COLO320 DM cell line^27^ was used to conduct live-cell imaging. Before cell seeding, 96-well glass-bottom plates were coated using poly-D-lysine. The medium was replenished with FluoroBrite DMEM (Gibco, A1896701), with 10% FBS, 1x GlutaMax and prolong live antifade reagent diluted in 1:200 ratio (Invitrogen, P36975) 30 minutes prior to imaging. The 96-well plate was fitted onto a top stage incubator (Okolab) and imaged on a Leica DMi8 widefield microscope using a x63 oil objective. Temperature (37°C), humidity and CO2 (5%) were stably maintained throughout the experiment.

### Micronuclei-harbouring cell fitness analysis via live-cell imaging

COLO 320DM-Tg19 cells were seeded in a coated well of a 96-well glass-bottom plate and treated with 50 µM hydroxyurea to enrich micronuclei formation. After 24 hours, hydroxyurea was removed and replaced with FluoroBrite DMEM (Gibco, A1896701), with 10% FBS, 1x GlutaMax and prolong live antifade reagent diluted in 1:200 ratio (Invitrogen, P36975). Z-stack images were acquired every 3 hours for a total of 60 to 72 hours, and images were processed using Small Volume Computational Clearing. Maximum Intensity projections were made on cropped frames of daughter cells immediately after initial division and for which at least one daughter cell contained a micronucleus. The micronucleus amplicon status (presence or absence of ecDNA⁺ or ecDNA-negative micronuclei) and micronucleus size (denoted as large if diameter is greater than 2 µm) of each daughter cell were recorded. The fate of daughter cells was manually tracked and assigned to the following three categories: continued division, cell death, or remaining undivided for the 72h imaging period. If the daughter cell divided, the timepoint at which the next division occurred was recorded to quantify time to next division (T2D). Cropped maximum intensity projections were imported into Aivia (v.12.0.0). Outlines were manually drawn around primary nuclei and their corresponding micronuclei. A machine learning-based pixel classifier and recipe were used to identify TetR-mNeonGreen foci by which ecDNA copy number in the primary nuclei and micronuclei were quantified. EcDNA status of micronuclei was classified as high or low relative to the median ecDNA content within micronuclei among all micronuclei.

### Microscopy, image acquisition and analysis

All images were acquired using a Leica Stellaris 8 confocal microscope at the Advanced Light Microscopy Technology Platform (Max Delbrück Center for Molecular Medicine). The system was equipped with an HC PL APO CS2 63×/1.40 OIL objective lens and used an immersion oil medium with a refractive index of 1.5185.

For imaging individual anaphase cells, the acquisition settings were as follows: 512 × 512 pixels frame size, 6× zoom, 0.79 µs pixel dwell time, and a final pixel size of 0.06 µm × 0.06 µm. Z-sampling was set to 0.23 µm to capture the entire cell in three dimensions.

For groups of interphase cells, images were acquired with a 2048 × 2048 pixels frame size, 2× zoom, 3.16 µs pixel dwell time, and a final pixel size of 0.05 µm × 0.05 µm.

Fluorophores were excited using diode laser lines at 561 nm (Alexa Fluor 568), 488 nm (Alexa Fluor 488), and 405 nm (Hoechst). Fluorescence emissions were detected at 566–620 nm (Alexa Fluor 568), 500–550 nm (Alexa 488), and 420–480 nm (Hoechst) using a pinhole set to 1 Airy unit (AU).

To ensure comparability across samples, all images were acquired under identical instrument settings.

The “Find Maxima” function was employed to quantify extrachromosomal DNA (ecDNA) copy numbers in FISH images. In anaphase cells, lagging DNA was defined as chromosomal material visibly separated from the primary segregating chromosomes by a minimum distance of 0.3 μm across the Z-stack, consistent with the lateral resolution limit of the imaging system. Images for Extended Fig 3b-d 3f, / Extended Fig. 8b were taken by a ZEISS LSM 880 inverted confocal microscope using ZEN (black v.2.3), and images for Fig 2bm, 2i-j, Extended Fig. 2d-f, 3h-i, Extended Fig. 8h-i were taken by a Leica DMi8 widefield microscope by Las X software (v.3.8.2.27713) using a ×63 oil objective. Z-stacks were taken for each field of view with image pre-processed using Small Volume Computational Clearing, and a max projection was performed by ImageJ (v.1.53t) before proceeding for further image analysis. Individual splitted channel images were semi-automatically segmented and analyzed by CellProfiler (v.4.2.1). Micronuclei were scored manually and defined as round structures separated from the primary nucleus, with a diameter up to one third of the size of the main nucleus, as previously described^25,39^.

Colocalization analysis was done using Huygens software (deconvolution, v24.10), FIJI (tile stitching, v2.14) and Imaris (colocalization analysis, v10.2).

### Nanopore-based replication fork dynamics

CHP212-PIP-FUCCI cells were seeded into T-300 flasks without Hygromycin B. At 70% confluency, cells were treated with or without 80 μM hydroxyurea for 24h. CHP212-PIP-FUCCI cells were then successively treated with 50 μM EdU for 3 minutes, 50 μM BrdU for 6 minutes, 100 μM Thymidine for 20 minutes, with 3 PBS washes in between each step. 80 μM hydroxyurea was maintained during the whole time, if treated. Cells were trypsinized, washed with PBS and sorted for S-phase (mCherry+, mVenus−) using a BD FACSAria Fusion. Sorted cells were pelleted, flash frozen in liquid nitrogen and stored at -80°C until DNA extraction. High molecular weight DNA was extracted using the NEB Monarch HMW DNA Extraction Kit for Tissue (NEB, T3060). DNA libraries for nanopore sequencing were prepared using the Ultra-Long DNA sequencing KIT (ONT, SQK-ULK001) according to the manufacturer’s instructions. Libraries were sequenced on R9.4.1 flow cells on the MinION platform.

Reads were basecalled using Guppy (v6.4.6, dna_r9.4.1_450bps_hac model, ONT) and aligned to hg38 using minimap2^40^ (v2.30). We used DNAscent (v3.1.2)^41^ to investigate replication fork dynamics. We applied the DNAscent detect function to detect EdU and BrdU incorporation on nanopore reads in batches of around 4000 reads. The forkSense function was then used to classify replication forks based on the order EdU and BrdU incorporation.

To determine replication fork speeds, we only retained complete forks (forks running off the end of a read were excluded) from the forkSense output. We then overlapped the remaining forks with the Decoil amplicon reconstruction for CHP212 and classified them as either circular (overlapping with ecDNA regions) or linear (overlapping with any other region). To compare ecDNA forks with linear forks of similar chromatin context, we further filtered forks which overlapped with active regions, by H3K27ac ChIPseq peaks. Bed files for peak calls were taken from^42^ and downloaded from GEO (GSE90683) and converted from hg19 to hg3838 using the UCSC liftover tool (https://github.com/ucscGenomeBrowser/kent). Fork speeds were calculated by dividing the length of each detected fork by the treatment time beginning with the start EdU treatment until the start of the thymidine chase out (Untreated: 15 min; HU: 16.4 min).

### Laser-microdissection and Whole Genome Amplification (WGA) of single (micro-)nuclei

TR-14 cells were directly grown on Poly-D-Lysine coated (Gibco, A3890401) PEN-membrane metal frame slides (Leica, 11600289). Immunofluorescence was performed as described above. Slides were stored in 1x PHEM in the dark at 4°C until dissection. Single micronuclei, pooled micronuclei (5-10 micronuclei) and single primary nuclei were manually visualised and identified on a Leica microdissection microscope (LMD7 or LMD7000, Leica microsystems). We collected primary nuclei and micronuclei from independent cells and primary and micronuclei from the same cell in a subset of samples (Extended Fig. 9d). Shapes were collected into adhesive PCR caps (MicroDissect GmbH, MDCA8W1K20). WGA was performed as in the ImmunoGAM method, with minor modification^43,44^. Sequencing libraries were prepared using the NEBNext Ultra II FS (New England Biolabs, E7805) kit according to the manufacturer’s protocol with reduced reaction volumes (1/4). Samples were barcoded for Illumina sequencing using the NEBNext Multiplex Oligos for Illumina kit (New England Biolabs, E7874) and sequenced on a NextSeq2000 sequencer (Illumina) or Novaseq X (Illumina) in 2×150-nt or 2 x 100-nt paired-end mode.

### Amplicon reconstruction in TR-14

Four distinct ecDNA elements were reconstructed for TR14 using Decoil^33^ 1.1.2 with parameters –min-vaf 0.1 –min-cov-alt 10 –min-cov 8 –fragmentmin-cov 10 –fragment-min***-***size 1000 –filter-score 35 or –min-vaf 0.01 –min-cov-alt 10 –min-cov 10 –max-explog-threshold 0.01 –fragment-min-cov 10 –fragment-min-size 500, with the reference genome GRCh38/hg38 and annotation GENCODE V42, (as in^33^). Additionally, a de novo assembly contig containing CDK4 for TR14 was assembled using Shasta^45^ (v0.6.0) with adjusted parameters minReadLength = 10000, k = 14, consensus caller Bayesian:SimpleBayesianConsensusCaller-10.csv. The contig was aligned to GRCh38/hg38 using minimap2^40^ with parameters -x asm10. From the PAF alignment the overlapping genomic coordinates were extracted as BED file. Using the genomic regions of all amplicons found in TR14 a BED file and a custom reference genome was computed, which includes (1) a masked GRCh38/hg38 genome of the regions overlapping the amplicons and (2) the FASTA sequence of all the ecDNA elements as additional contigs.

### Identification of micronucleated sequences

Reads were quality and adapter-trimmed using Trim Galore^46^ (v0.6.10) and aligned to the human reference assembly GRCh38 with added EBV contig (NC_007605.1) using BWA-MEM^47^ (v 0.7.18) with default parameters. Duplicate reads were removed using Picard^48^ (v3.3.0) MarkDuplicates. We collected all samples over a total of 2 experiments which were sequenced on different flowcells. Samples from the run with higher sequencing depth were downsampled to match the median number unique reads from the other run using samtools view with the -s parameter. BigWig files were generated using Deeptools2^49^ (v3.5.5) bamCoverage with a binsize of 25kb and CPM normalization.

### Candidate window identification

For each sample, we identify candidate windows, which represent genomic regions covered by reads significantly over background, detected by performing the following 4-steps: (1) The genome was divided into 25kb bins and the number of reads and number of bases covered by at least one read were counted with bedtools coverage. (2) For each sample, we fitted a negative binomial distribution on the per-bin read counts, similar to^50^. Distribution parameters were estimated using Maximum Likelihood Estimation as implemented in the fitdistrplus^51^ (v1.2.1) R-Package. The threshold RC_th_ for per-bin read counts was derived from the first read count value of the cumulative mass function, where the probability of it being larger than threshold RC_th_ was <0.001, as defined as >1-P, where P denotes the probability of a value being less or equal than threshold RC_th_. (3) To remove bins with focal read accumulation, thus being bins with high read counts but low coverage, we further sorted all bins based on the percentage of bases covered. We defined a coverage threshold C_th_ as the 99th percentile. (4) Only bins with a read count > RC_th_ and coverage > C_th_ where retained as candidate windows.

To identify enriched sequences under the assumption that DNA sequestered into micronuclei, which are detectable by light microscopy, derive from chromosomes, chromosomal fragments or other longer chromosomal units (such as ecDNA) of a certain length, we further refine the candidate window set by removal of solidary windows, which adhere to the candidate window definition as described above, but do not have any adjacent candidate windows or are of small size. With this approach we limit the smallest detectable unit size, but make sure to further remove regions, which are most likely derived from amplification and/or mapping noise.

We pivot through each candidate window and search for adjacent candidate windows up and downstream within a radius r. A radius of r=4 was used in this study. If a candidate window has at least one other candidate window upstream and downstream within the radius r, this candidate window is further classified as a body segment, while candidate windows with only at least one other candidate window within the radius r on either side, are classified as edge segment. If a candidate window does not have any other candidate windows within its radius, this window is removed from analysis. We further align edge and body segments into closed segments, where body segments must be flanked by edge segments to be retained, thus making an edge-body-edge alignment the smallest detectable unit, which for this experiment is a fragment size of 75kb. If an edge segment is flanked by another edge segment, both windows are removed from further analysis.

### Copy number calling

Copy number profiles for each sample were computed similarly to CNVkit^52^ using a custom python script utilizing the Pysam API (https://github.com/pysam-developers/pysam). First, the genome was tiled into 500kb bins. For each bin, per base coverages were summed up and divided by the bin size to get a per bin mean coverage, which was then log2 transformed.

Visualization and segmentation of copy number profiles were computed using a custom R script (https://github.com/henssen-lab/ecMN/tree/main/copy_number). First, only canonical chromosomes (autosomes and sex chromosomes) were retained and telomeric and centromeric regions filtered out. GC content for each bin was calculated using the R package Biostrings^53^. We performed GC correction for the log2 per bin mean coverage values using the loess function from the stats R-package with a smoothing span of 0.3 and performed median centering based on all autosomal bins. Segmentation was performed using circular binary segmentation, as implemented in the DNAcopy^54^ R-package. After smoothing and outlier removal using the smooth.CNA function, segments were called using the segment function with following parameters: undo.splits = “sdundo”, undo.SD = 1.5. Log2 per bin coverages and segments were visualized with ggplot2^55^.

### (Micro-)nuclear sequence analysis

#### Quality control and filtering

Single and pooled micronuclei samples were retained for downstream analysis if they contained more than 20 called windows and >100000 uniquely mapping reads. We additionally added a second quality control layer, where we inspected the copy number profiles and excluded samples, which showed high degrees of background amplification. Primary nuclei samples were retained if they had more than 50 called windows.

#### EcDNA consensus sequence calling in single micronuclei

After visual inspection of called regions in single micronuclei, we observed an enrichment of sequences overlapping the amplicon regions, as described above. To identify the consensus region between enriched sequences in single micronuclei and ecDNA reconstructions for further quantification, we generated a set of all called windows from single micronuclei passing quality control. We retained each genomic region, which was called in at least 2 samples. We overlapped these regions with the custom reconstructions and were able to retain the majority of ecDNA regions in single micronuclei. The only region we could not capture was a short region in chromosome 1, whose length fell below the smallest detectable unit size of 75kb as described above. The consensus ecDNA regions were used for subsequent analysis.

#### Circular read enrichment estimation (log2 coverage)

To quantify read enrichment over the consensus ecDNA regions, we further normalized the absolute per-bin read counts to counts per million to account for different library size. To estimate the enrichment of circular reads, we divided the mean coverage of all consensus ecDNA regions by the winsorized mean coverage (top and bottom 5% values replaced) of all non-ecDNA regions, as implemented in the Winsorize function from the R-Package DescTools^56^ (v0.99.58). The coverage ratio was further log2 transformed.

To quantify the signal over each connected consensus ecDNA region over their flanking region, we obtained region-scaled coverage values using Deeptools2 computeMatrix scale-regions with parameters: -m 250000 -b 125000 -a 125000 -bs 25000 –missingDataAsZero – outFileNameMatrix.

#### Co-segregation and relative ecDNA fraction analysis

To assess ecDNA co-segregation heterogeneity, we counted the presence or absence for each amplicon in all single micronuclei and primary nuclei samples. For each sample, we overlapped all called windows with the amplicon reconstructions. An amplicon was deemed present, if it had at least 1 overlapping called window.

To evaluate the relative copy number differences between different amplicons within one sample we computed amplicon fractions within all circular reads. Read counts for each amplicon were first divided by its respective amplicon length to normalize counts for amplicon size. The length normalized counts for each amplicon were further divided by the sum of the length normalized counts of all amplicons to obtain a value, which represents the fraction of reads belonging to each respective amplicon within all circular reads.

#### Linear chromatin enrichment analysis

To identify chromosomal fragments, which are enriched in single micronuclei, we first accumulated per chromosome read counts, excluding all reads mapping to the amplicon regions, and normalised them to chromosome size. Enriched chromosomes have high read accumulation over background mapping. We therefore identified enriched chromosomes as extreme outliers, if the normalised read count was larger than: Q3 (3^rd^ quartile) + 10 * IQR (interquartile range). The fragment length of the enriched chromosome was determined by summing up the number of bins, where the per bin read count was larger than the 90^th^ percentile of all non-zero bins within the identified enriched chromosome.

#### Chromatin immunoprecipitation (ChIP)-sequencing

Hydroxyurea- and DMSO-treated COLO320DM cells were washed 2x with prewarmed 1X PBS. Cells were fixed by addition of 10mL ice-cold 1% FA in 10% FCS-RPMI-1640 and incubated for 10 min on a shaker. The fixation reaction was quenched by addition of 550uL ice-cold 2.5M Glycine in PBS. Cells were washed 1x with ice-cold 1X PBS before scraping down in 10 mL ice-cold 1X PBS. Fixed cells were centrifuged at 500 x g for 5 min at 4°C and resuspended in ice-cold 1X PBS for a total of 2 washes. Cells were pelleted and snap frozen in liquid nitrogen and stored at -80 C for no more than 1 week.

Cell pellets were resuspended in 5 mL ice-cold lysis buffer 1 (140 mM NaCl; 50 mM HEPES-KOH pH 7.5; 1 mM EDTA; 10% Glycerol (v/v); 0.5% IGEPAL CA-630 (v/v); 0.25% Triton X-100 (v/v); 1x Protease inhibitor (Roche, 04693132001)) and incubated on a roller at 4°C for 10 min. Lysed cells were centrifuged at 1357 x g for 5 min, resuspended in 4 mL lysis buffer 2 (20 mM NaCl; 10 mM Tris-HCl pH 8.0; 2 mM EDTA; 0.5M EGTA; 1x Protease inhibitor) and incubated on a roller for 10 min at room temperature. The lysate was centrifuged at 1357 x g for 5 min at 4°C and resuspended in 900 μL sonication buffer (100 mM NaCl; 10 mM Tris-HCl pH 8; 1 mM EDTA; 0.5 EGTA; 0.1% Na-deoxycholat (w/v); 0.5% N-Laroylsarcosine (w/v); 1x Protease inhibitor). Lysed chromatin was sheared to a fragment size of 200-500 base pairs (bp) on a Covaris S220 (PIP 140, DF 5, CPB 200) for 12 min. Sheared chromatin was clarified by addition of 1/10 volume 10% Triton X-100 (v/v) and centrifugation at maximum speed at 4°C.

Protein-DNA-complexes were immunoprecipitated by overnight incubation with the respective primary antibody on a rotator at 4°C. 10-15 μg total chromatin was used per sample. 30 μL Protein G beads (Invitrogen, 10003D) were washed 3x in 0.25% BSA in PBS (w/v) and added to each sample. Samples were incubated overnight on a rotator at 4°C.

Antibody-bead - complexes were washed in RIPA buffer (50 mM HEPES-KOH pH 7.5; 1mM EDTA; 1% IGEPAL CA-630 (v/v); 0.7% Na-deoxycholat (w/v); 500 mM LiCl; 1x Protease inhibitor) for a total of 7 times followed by 1 wash in TE buffer (10 mM Tris-HCl pH 8.0; 1mM EDTA; 50 mM NaCl; 1x Protease inhibitor). Beads were eluted in 200 μL Elution Buffer (50 mM Tris-HCl pH 8.0; 10 mM EDTA, 1% SDS (v/v)). 20 μL 5M NaCl and 5 μL Proteinase K (NEB, P8107S) were added to each sample and DNA was de-crosslinked at 65°C overnight, followed by RNA digestion with 2 uL RNase A (Invitrogen, R1253) at 37°C for 30 minutes. DNA was purified using the QIAquick PCR Purification Kit (Qiagen, 28104) according to the manufacturer’s instructions. ChIP DNA libraries were constructed using the NEBNext Ultra II DNA Library Prep Kit (NEB, E7645S) and barcoded for Illumina sequencing using the NEBNext Multiplex Oligos for Illumina kit (NEB, E7780S). Libraries were sequenced on a NovaSeq X Plus sequencer (Illumina) in 100 nt single-end mode.

#### ChIP-sequencing analysis

The raw reads were quality and adapter-trimmed using Trim Galore and aligned to the human reference assembly GRCh38 with an added EBV contig (NC_007605.1) using BWA-MEM with default parameters. Duplicate reads were removed using Picard MarkDuplicates. Library quality was assessed using strand cross-correlation metrics (RSC and NSC) from Phamtompeakqualtools^57,58^ (v1.2.2).

Using the deduplicated BAM file, read densities per 10 bp window were generated using DeepTools2 bamCoverage. Genomic bins overlapping the ENCODE DAC blacklist regions were filtered out using the parameters: --ignoreForNormalization chrX chrM NC_007605.1 --scaleFactor 10 --effectiveGenomeSize 2805636231 --exactScaling --extendReads 200 --normalizeUsing CPM and input-substracted using the bigwigCompare function.

#### RNA-sequencing

Three replicates of Hydroxyurea- and DMSO-treated COLO320DM cells were washed with 1x PBS, trypsinized (Gibco) and centrifuged at 300xg for 5 min. Total RNA was immediately extracted from the cell pellets using the RNeasy Mini Kit (Qiagen, 74104) following the manufacturer’s instructions. Genomic DNA was removed by DNase treatment using the RNase-Free DNase Set (Qiagen,79254). The mRNA library was constructed using the TruSeq Stranded mRNA Library Prep kit (Illumina) at the Berlin Institute of Health (BIH) Genomics Technology Platform. Libraries were sequenced on a NovaSeq X Plus sequencer (Illumina) in 100 nt single-end mode.

#### RNA-sequencing analysis

Reads were quality and adapter-trimmed using Trim Galore and aligned to the human reference assembly GRCh38 with an added EBV contig (NC_007605.1) using STAR^59^ (v2.7.11b) with parameters --twopassMode Basic --outSAMtype BAM SortedByCoordinate. Reads were counted per gene against GENCODE gene annotation (Release 47) using htseq^60^ (v2.0.5) htseq-count with parameters -r pos -s no. Counts were normalized for library size and composition using sizefactors from DEseq2^61^ (v1.38.3). Counts were z-score normalized for visualization using the scale function in R-base (v4.2.2). Differentially expressed genes between the untreated and treated samples were identified using DEseq2 and p-values were corrected for multiple testing using the Benjamini-Hochberg procedure. Gene set enrichment analysis was performed on differentially expressed genes with an adjusted p-value <0.05 against MSigDB hallmark gene sets using the GSEA function in Clusterprofiler^62^ (v4.6.2) and visualized with DOSE^63^ (v3.24.2).

#### Patient samples and clinical data access

This study comprised the analyses of tumour and blood samples of patients diagnosed with neuroblastoma between 1991 and 2022. Patients were registered and treated according to the trial protocols of the German Society of Pediatric Oncology and Hematology (GPOH). This study was conducted in accordance with the World Medical Association Declaration of Helsinki (2013) and good clinical practice; informed consent was obtained from all patients or their guardians. The collection and use of patient specimens and clinical data was approved by the institutional review boards of Charité-Universitätsmedizin Berlin and the Medical Faculty, University of Cologne. Specimens and clinical data were archived and made available by Charité-Universitätsmedizin Berlin or the National Neuroblastoma Biobank and Neuroblastoma Trial Registry (University Children’s Hospital Cologne) of the GPOH. The *MYCN* copy number was determined as a routine diagnostic method using FISH. DNA and total RNA were isolated from tumour samples with at least 60% tumour cell content as evaluated by a pathologist.

#### Clinical data analysis

Patients were registered in the GPOH trials NB90^64^, NB97^65^, NB2004 (NCT00410631) and NB2016-registry. All protocols were approved by the ethics committees of the Universities of Cologne as leading institutions (see supplemental table 2 for approval numbers), and approval was confirmed by the local ethics committees. Informed consent was obtained from the participants at the local treatment site and was a prerequisite to register in the respective trials. Trial participation was a requirement for inclusion in this analysis. For the purpose of this analysis, follow-up information was evaluated for matching tumour samples of a respective patient. The endpoints were overall and event-free survival. Overall survival (OS) was defined as death of any cause and event-free survival (EFS) was defined as disease progression or relapse, or death of any cause. Patients without event were censored at last follow-up. OS and EFS were calculated using the Kaplan-Meier estimator^66^. The log rank test was used for comparison. A Cox proportional hazards regression was conducted for ecDNA+ micronuclei percentage. Statistical analysis was conducted using SPSS 29 (Armonk, New York, U.S.). Despite availability, exclusively clinical data of samples collected at first diagnoses before the start of therapy were included to eliminate a potential bias due to therapy-related changes of tumour genetics.

### Modelling of ecDNA segregation patterns

#### Estimation of ecDNA mis-segregation probability

To estimate the mis-segregation probability of ecDNA in a patient or cell line, we adapted a binomial model of micronucleation^67^. According to this model, the frequency of cells without micronuclei is given by 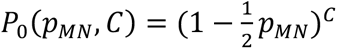, where pMN is the mis-segregation probability of an ecDNA element during cell division and C is the amount of ecDNA elements in the cells. Per patient and cell line where we have more than five cells imaged, we then use the average observed ecDNA copy number by FISH per cell and frequency of non-micronucleated cells to calculate p_MN_.

#### Biased Random Segregation Model

We developed a model to simulate ecDNA segregation during cell division. A single cell begins with a random ecDNA copy number 𝑛, sampled from a uniform distribution. Prior to cell division, ecDNA is amplified to 2𝑛 copies, which are then partitioned between two daughter cells. For cells undergoing random segregation, ecDNA partitioning follows a binomial distribution 𝐵(2𝑛, 𝑝_2_), where 𝑝_2_ = 0.5. In contrast, for cells exhibiting biased random segregation, partitioning follows a binomial distribution 𝐵(2𝑛, 𝑝_1_), where 𝑝_1_ < 0.5 reflects asymmetric segregation. The probability of biased segregation is denoted as 𝑞, while 1 − 𝑞 represents the probability of random segregation. The ecDNA copy numbers in the two daughter cells are denoted as 𝑛_1_ and 𝑛_2_ = 2𝑛 – 𝑛_1_, respectively. The ecDNA fractions 𝑓_1_ and 𝑓_2_ of the two daughter cells can be calculated as: 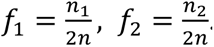. By repeating this process, we generate the distribution of ecDNA fractions across daughter cells.

#### Reconstructing ecDNA Segregation Distribution under Finite Sampling

We first simulated ecDNA segregation in 10^7^ cells to establish the “true” or “full” distribution. We then sampled subsets of cells at sample sizes of 10, 25, 50, 100, and 500, repeating each sampling 10^5^ times. To quantify the deviation between sampled and true distributions, we calculated the Kolmogorov distances.

#### Parameter Estimation Using Approximate Bayesian Computation (ABC)

We applied Approximate Bayesian Computation (ABC) to infer the parameters 𝑝_1_ and 𝑞 in the model. For the simulated data, 50 pairs of daughter cells were randomly drawn using predefined parameters. During inference, 𝑝_1_and 𝑞 were sampled from prior uniform distributions: 𝑝_1_∼𝑈(0, 0.5) and 𝑞∼𝑈(0, 1). 50 parent cells were randomly selected, with their ecDNA copy numbers drawn from a uniform distribution 𝑈(10, 100). For each set of parameters, we ran the model to obtain the ecDNA segregation fraction distribution. To compare the similarity between simulated and target distributions, we computed the Wasserstein distance. A threshold was applied to determine acceptance, generating the posterior distributions of 𝑝_1_ and 𝑞. For experimental data, the same procedure was followed, with the sample size adjusted according to the specific experimental conditions.

### Statistical analysis

All experiments were performed a minimum of two times with a minimum of three independent measurements unless otherwise stated. All statistical analysis was performed with R 3.6, R 4.0 or Python 3.7. All data are represented as mean ± standard error and tested for normal distribution. Statistical significance was defined as *, *P* < 0.05; **, *P* < 0.01, ***, *P* < 0.001, ****, P < 0.0001. Normality was assessed using the Shapiro-Wilk test.

## Data availability

Sequencing data produced in this study will be deposited in the European Nucleotide Archive (https://www.ebi.ac.uk/ena/browser/submit). Custom TR14 amplicon reconstructions and FASTA will be available under github.com/henssen-lab. The data that support the findings of this study are available from the corresponding author upon request.

## Code availability

Scripts for data analysis within the scope of this publication will be available under https://github.com/henssen-lab/ecMN.

## AI assisted copy editing

All text was written by the authors. Parts of the text were copy edited using ChatGPT to improve readability and clarity.

## Materials & Correspondence

Correspondence and requests for materials should be addressed to

## Author Contributions

L.B., R.X., J.T., A.G., A.H., I.T.W., and S.Z., contributed to the study design and collection and interpretation of the data and wrote the manuscript. F.T, M.P, K.P., O.S., J.S., Q.Y., D.P., M.I., A.K., S.H., M.G., A.S.H., M.F., F.B., V.R.L., C.D., X.Y., D.G., S.K., D.G., M.R. N.W., G.M. N.W., performed the experiments, analyzed data and reviewed this manuscript. J.T., I.T.L.W., S.Z., performed the experiments, analyzed and interpreted the data, and edited the manuscript. D.T., A.L., F.D. and M.F. provided data. B.P. and A.H. established essential methods. M.S. analysed the clinical data. R.L., G.B. Y.L., V.B., G.W., R.M., S.P., B.S., A.P., R.P.K., W.H., and B.W. contributed to study design. A.G.H., P.S.M., H.Y.C. led the study design, to which all authors contributed.

## Acknowledgements

This work was delivered as part of the eDyNAmiC team supported by the Cancer Grand Challenges partnership funded by Cancer Research UK and led by P.S.M. (CGCATF-2021/100017) (A.G.H.), (CGCATF-2021/100012) (P.S.M. and H.Y.C.), (CGCATF-2021/100020) (B.W and W.H), (CGCATF-2021/100025) (V.B.), and the National Cancer Institute (OT2CA278644) (A.G.H.), (OT2CA278688) (P.S.M. and H.Y.C.), (OT2CA278670) (B.W and W.H), (OT2CA278635) (V.B.), A.G.H. is supported by the *Deutsche Krebshilfe* (German Cancer Aid) Mildred Scheel Professorship program – 70114107. This project has received funding from the European Research Council (ERC) under the European Union’s Horizon 2020 research and innovation programme (grant agreement No. 949172). This project was supported by Cancer Research UK and the National Institute of Health (398299703, the eDynamic Cancer Grand Challenge). Research in the G.B. laboratory is supported by the UK Dementia Research Institute, which receives contributions from the Medical Research Council, UK and the CHDI Foundation, USA. Research at TINS is supported by Romanian PNRR-HUNTGEN-C9-I8 - Contract no. 760114/2023, financed by the European Union – NextGenerationEU. R.X. was supported by the Deutsche Forschungsgemeinschaft (DFG, German Research Foundation) funded Research Training Group (RTG) 2424/CompCancer – project number: 377984878. Work by D.P. in the Andrea Venturas. group was supported by grants from NIH-NCI (P30 CA008748) and (R01 CA282913 to AV), the American Cancer Society Discovery Boost grant (AV), the The Mark Foundation for Cancer Research ASPIRE grant (AV) and by a rapid response grant from the Functional Genomics Initiative (AV). D.P. was supported by an AIRC fellowship for Abroad. The authors want to thank Steven Whittaker for providing important conceptual input. B.W. is supported by a Barts Charity Lectureship (grant no. MGU045) and a UKRI Future Leaders Fellowship (grant no. MR/V02342X/1). W.H is also supported by NNSF General Program (grant no. 3217024. We thank B. Hero, H. Düren, N. Hemstedt of the Neuroblastoma Biobank and Neuroblastoma Trial Registry (University Children’s Hospital Cologne) of the German Society of Pediatric Oncology and Hematology (GPOH) for providing samples and clinical data. We thank Sabine Taschner at the St. Anna Kinderkrebsforschungsinstitut for providing cell lines. We thank Andrea Ventura at MSKCC for the establishment and data of the engineered mouse model. Computation has been performed on the HPC for Research cluster of the Berlin Institute of Health. R.P.K. is a Visiting Professor funded by the Stiftung Charité. We thank Hans-Peter Rahn at the MDC Flow Cytometry for support with FACS. Images were acquired at the Advanced Light Microscopy facility of the Max Delbrück Center for Molecular Medicine and the Systems Imaging Facility of the Berlin Institute for Medical Systems Biology.

