## Extended Data for "Extrachromosomal DNA micronucleation constrains tumour fitness and improves patient survival"

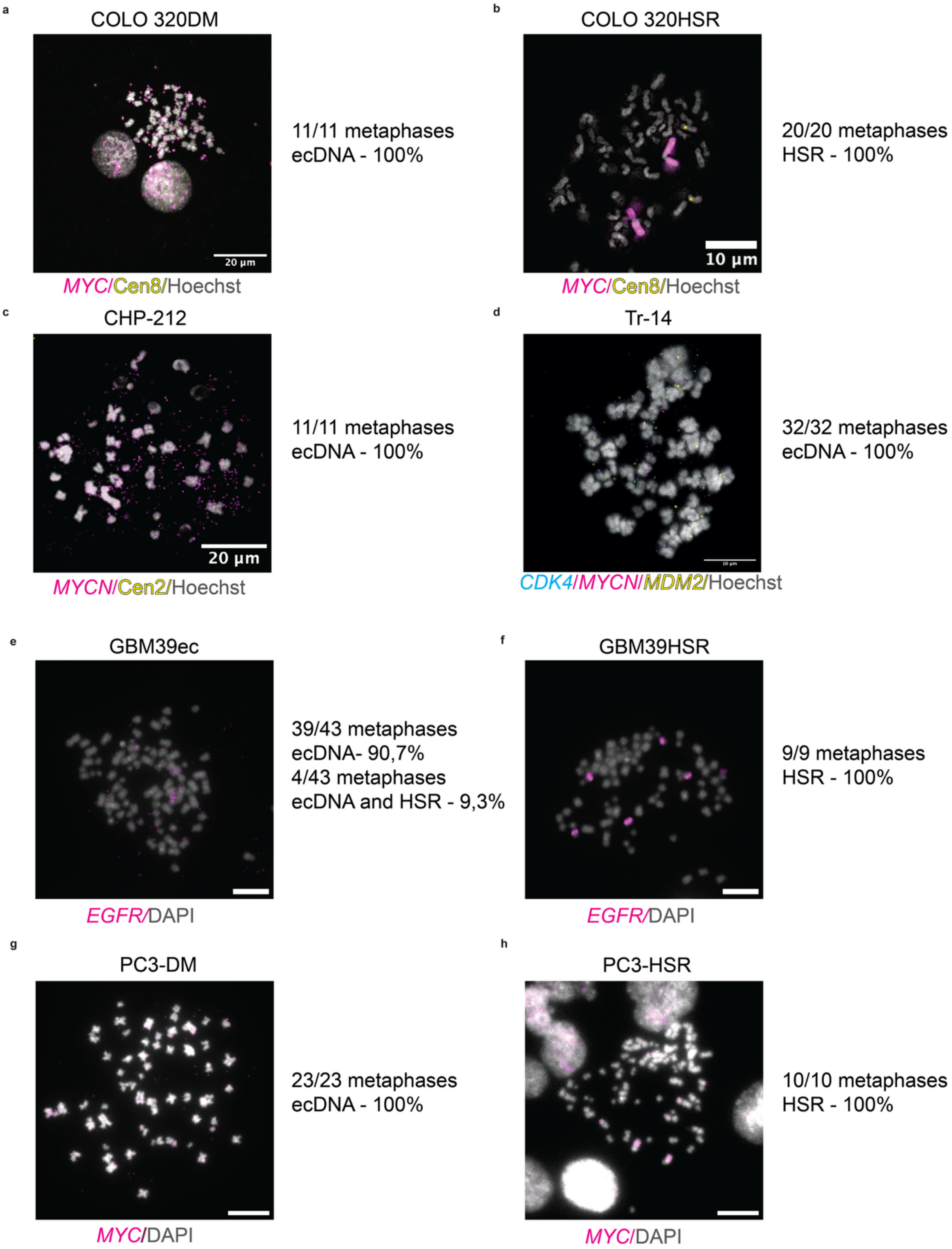


**Extended Fig. 1: Confirmation of amplification status of the major cell line models used in this study**

a. Exemplary metaphase spread and quantification of amplification status at early (p= 5) passage of the COLO 320DM single cell clone. Scale bar: 20 µm.

b. Exemplary metaphase spread and quantification of amplification status of the COLO 320HSR cell line. Scale bar: 10 µm.

c. Exemplary metaphase spread and quantification of amplification status of the CHP-212 cell line. Scale bar: 20 µm.

d. Exemplary metaphase spread and quantification of amplification status of the Tr-14 cell line. Scale bar: 10 µm.

e. Exemplary metaphase spread of the GBM39ec cell line. Scale bar: 10 µm.

f. Exemplary metaphase spread of the GBM39HSR cell line. Scale bar: 10 µm.

g. Exemplary metaphase spread of the PC3-DM cell line. Scale bar: 10 µm.

h. Exemplary metaphase spread of the PC3-HSR cell line. Scale bar: 10 µm.

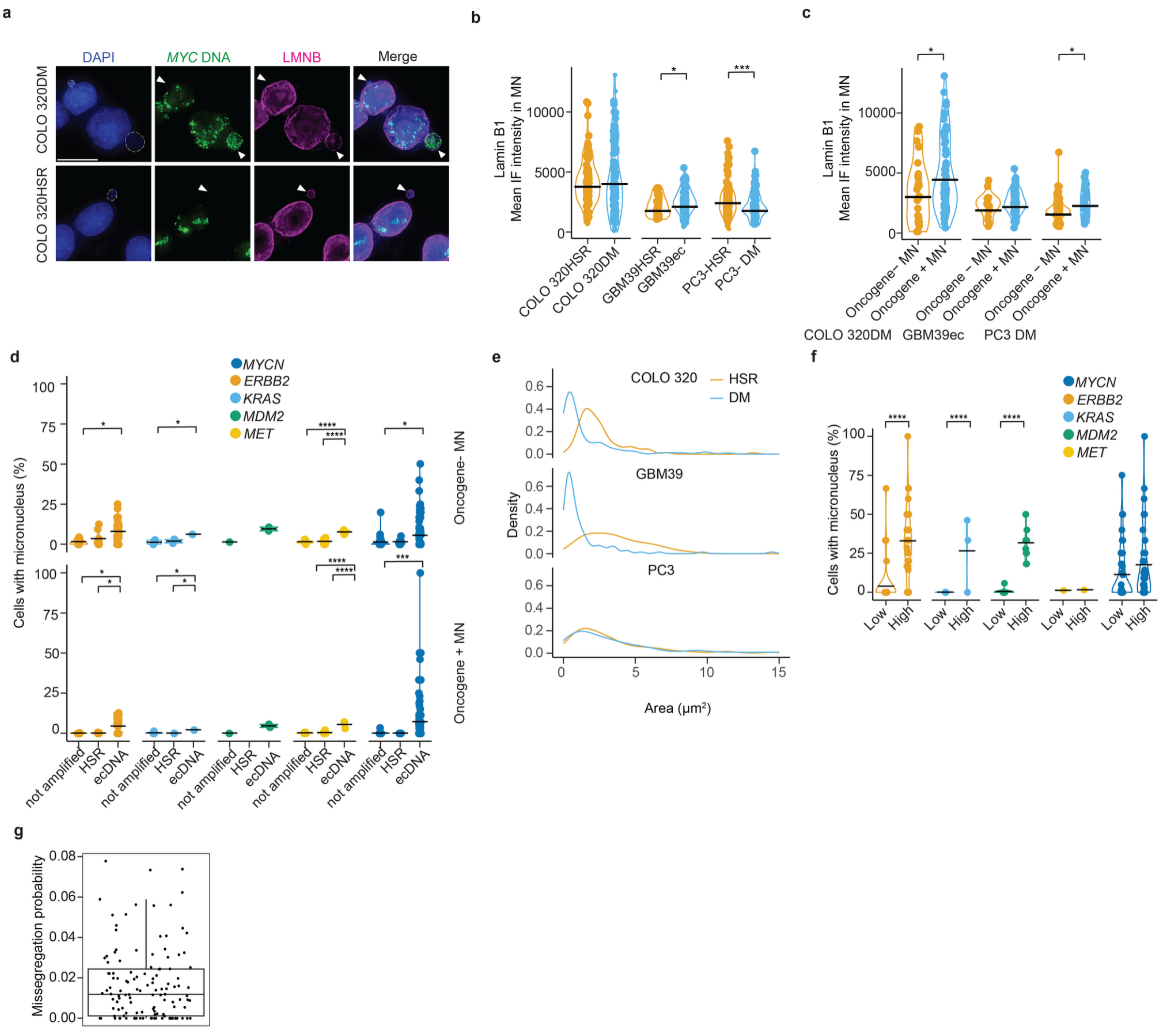

**Extended Fig. 2: Cancer cells with high ecDNA content are prone to micronucleation irrespective of the amplified gene**

a. Exemplary photomicrographs of COLO 320DM and COLO 320HSR cells with micronucleus (white arrowhead and dash) stained for *MYC* and lamin B1 by ImmunoFISH. Scale bar: 10 µm

b. Mean lamin B1 staining intensity in micronuclei in three near-isogenic cell line pairs (n= 32-107, Student's t-test). Each dot represents one micronucleus.

c. Mean lamin B1 staining intensity in ecDNA$^+$ MN and ecDNA$^-$ MN determined by FISH in three near-isogenic cell line pairs (n= 22-84, Student's t-test). Each dot represents one micronucleus.

d. Fraction of cells with micronucleus in patient tumor samples stratified by amplified gene and oncogene status in MN (Student's t-test).

e. Density plot displaying micronuclei size distribution in three near-isogenic cell line pairs: COLO 320DM (n=103), COLO 320HSR (n=100), GBM39EC (n=100), GBM39HSR (n=33), PC3-DM (n=103) and PC3-HSR (n=102).

f. Fraction of cells with micronuclei and low (bottom 30%) vs. high (top 30%) ecDNA content measured in patient tumor samples stratified by amplified gene (j) (Student's t-test). Black lines indicate the mean.

g. Mis-segregation probabilities (the chance that at least one ecDNA micronucleates during mitosis) across 8 cell lines (COLO 320DM, CHP-212, STA-NB10DM, Tr-14, UKF-NB6, Lan5, SMS-KAN,KP-N-YN) and 119 patient samples.

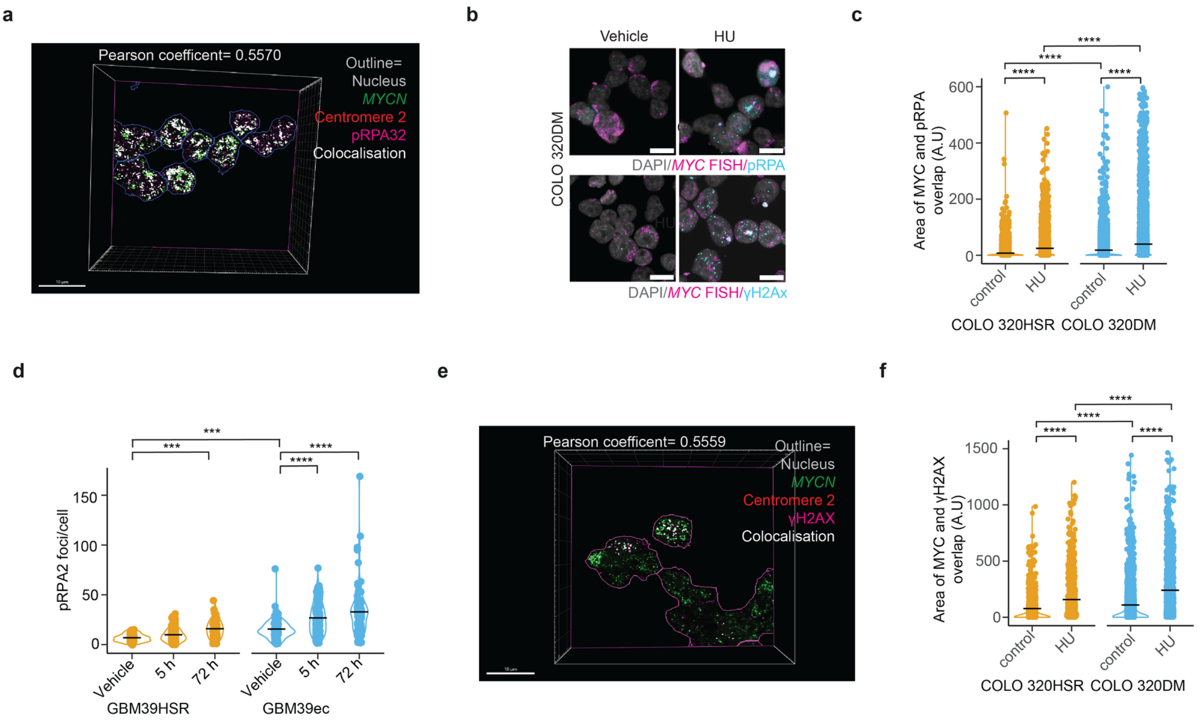


**Extended Fig. 3: Replication stress and DNA damage induction on ecDNA**

a. Representative 3D reconstruction of fixed untreated CHP212 cells, that contain *MYCN* ecDNA, stained for Hoechst (grey outline), *MYCN* (green, FISH), centromere 2 (red, FISH) and pRPA32 (magenta, immunofluorescence). Areas where *MYCN* ecDNA (green) and pRPA32 (S4/S8) (magenta) colocalise, were computed using Imaris (v10.2) and are depicted in white. Scale bar: 10µm.

b. Exemplary photomicrographs of fixed COLO 320DM cells in the presence of HU (50 µM, 72h) or vehicle, stained for *MYC* (magenta, FISH) and pRPA32 (blue, immunofluorescence) (top row) or *MYC* (magenta, FISH) and γH2AX (blue, immunofluorescence) (bottom row). Scale bar:10µm.

c. Quantification of colocalization between pRPA2 and *MYC* FISH in COLO 320DM and COLO 320HSR to indicate replication stress on the amplicon. (n = 3602–8424; heteroscedasticity-robust two-way ANOVA). Each dot denotes one cell.

d. Quantification of pRPA2 foci per cell in isogenic glioblastoma cell lines at different timepoints of 100µM HU treatment (n= 68-175, Two-way ANOVA). Each dot represents one cell.

e. Representative 3D reconstruction of ImmunoFISH stained cells of *MYCN* and γH2AX in untreated CHP-212 cells. Areas of colocalisation are indicated in white. Colocalisation was calculated between Channel 2 (*MYCN*, green) and Channel 4 (γH2AX, magenta).

f. Quantification of γH2AX foci colocalised with *MYC* in an isogenic colon cancer cell line pair after 50 µM HU treatment for 3 days (n = 596- 1648; heteroscedasticity-robust two-way ANOVA). Each dot denotes one cell.

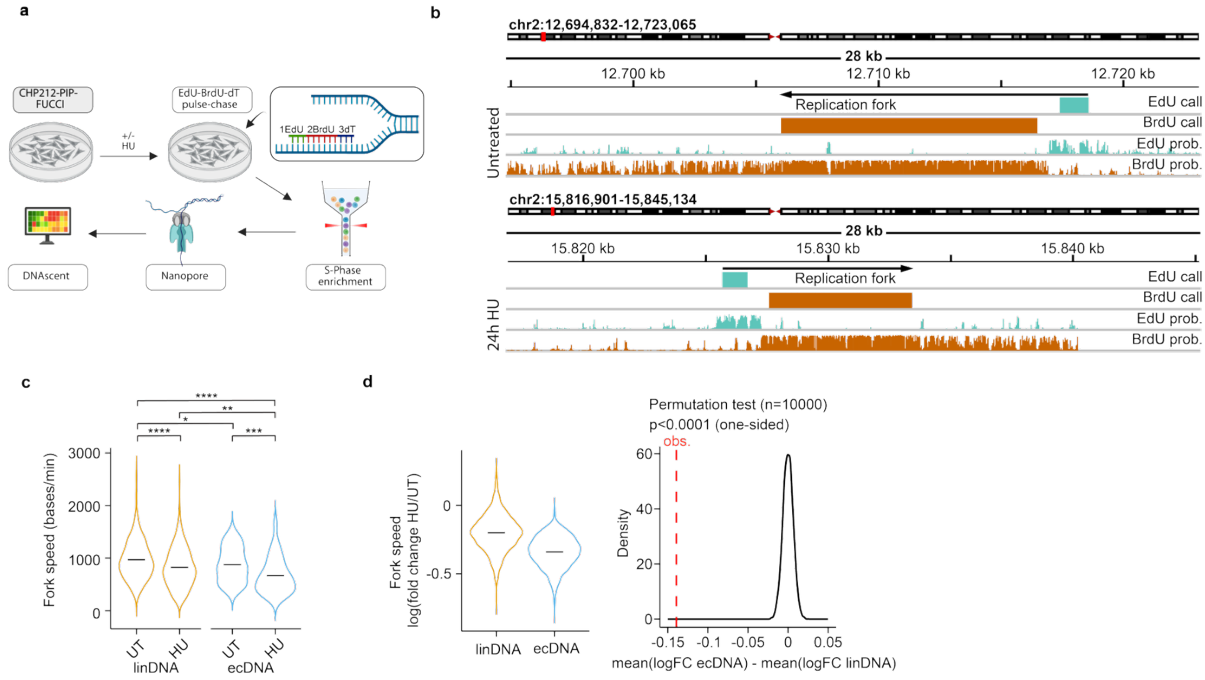


**Extended Fig. 4: Hydroxyurea induces replication fork slowing on ecDNA.**

a. Schematic of the experimental design. CHP212-PIP-FUCCI cells were incubated in the presence or absence of HU (80 µM) for 24h and subsequently treated with EdU, BrdU and dT. Cells were enriched for S-phase using FACS and sequenced using Nanopore (ONT). DNAscent[1] was used to infer replication fork dynamics.

b. Representative IGV snapshot of EdU and BrdU tracks on ecDNA in CHP212-PIP-FUCCI cells in the presence or absence of HU (80 µM, 24h). The top 2 rows denote EdU and BrdU calls from DNAscent forksense. The bottom 2 rows denote the EdU and BrdU probability called by DNAscent detection. Arrows show the movement direction of the exemplary replication forks.

c. Quantification of replication fork speed, as computed by replication fork length divided by treatment time (bases/min) on linear DNA (linDNA), or ecDNA in the presence or absence of HU (80 µM, 24h).(UT-linDNA, n = 598; HU-linDNA, n = 535; UT-ecDNA, n = 73; HU-ecDNA, n = 63; Welch's t-test).

d. Low dose hydroxyurea (80 µM, 24h) has a differential effect on ecDNA. Left: Log foldchanges of HU-treated vs. untreated forkspeeds on linear DNA (linDNA) and ecDNA, as computed by subsampling 30 fork speeds per treatment and representad as log foldchange between the median fork speeds between HU treatment and untreated fork speeds (n=1000 repetitions per codition). Right: Comparison of the differential mean fork speed log foldchanges between linear and ecDNA (p<0.0001, permutations= 10000, permutation test, one sided). Red line indicates observed mean differential log foldchange from the left panel.

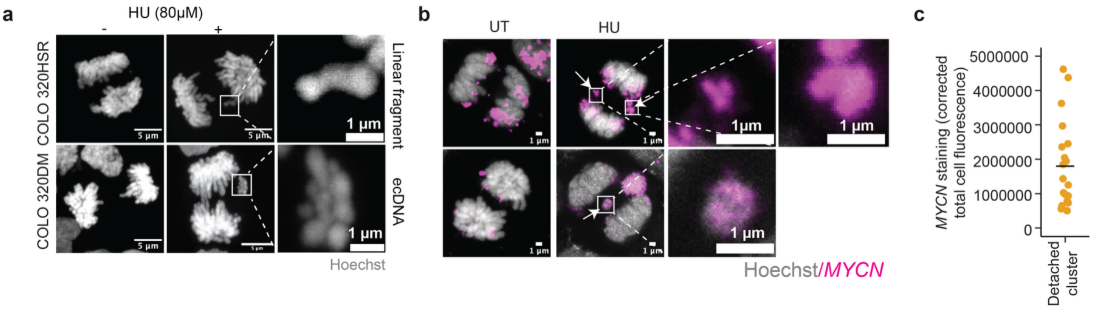


**Extended Fig. 5: Detached DNA clusters in CHP-212 are positive for amplicon staining**

a. Exemplary photomicrographs of representative fixed COLO 320DM (contains *MYC* as ecDNA, bottom) and COLO 320HSR (contains *MYC* as HSR, top) cells in anaphase that were incubated in the presence or absence of HU (80µM) for 3 days and stained with Hoechst. Scale bar: 5 µm and 1 µm (zoom). White box indicates enlarged region

b. Exemplary photomicrographs of fixed CHP212 cells during anaphase without (first column) and with HU treatment (second column) (80 µM, 24h), stained for Hoechst (grey) and *MYCN F*ISH (magenta). Arrows indicate detached DNA cluster. White boxes indicate enlarged regions. Scale bar: 1 µm.

c. Quantification of the corrected total cell fluorescence of *MYCN* FISH signal on detached clusters during anaphase in fixed CHP212 cells after HU treatment (80 µM, 24h, n=100 anaphases).

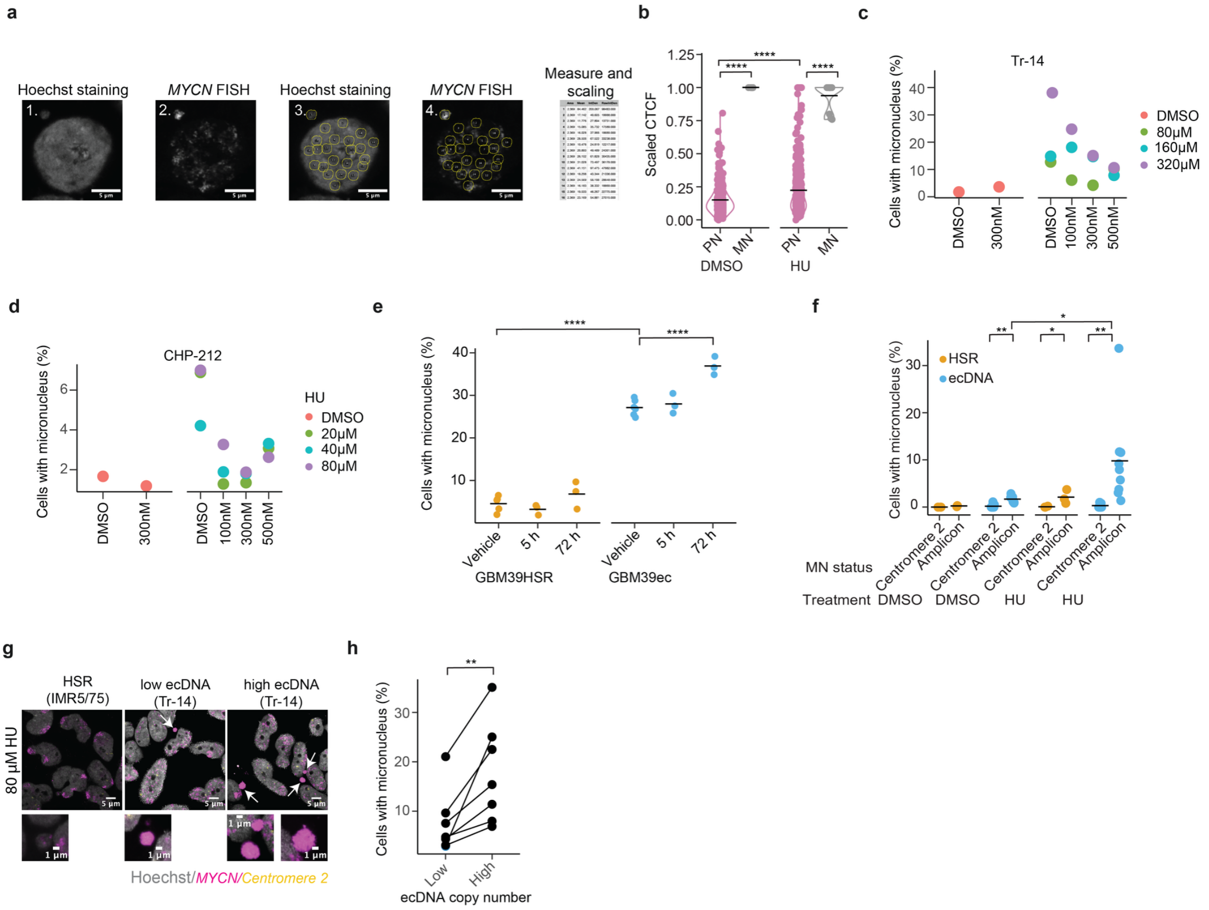


**Extended Fig. 6: Induction of replication stress induces ecDNA micronucleation**

a Schematic setup of the image analysis approach, quantifying fluorescent intensity and calculating the corrected total cell fluorescence in the micronuclei and primary nuclei (same size area). Scaling was done to account for differential background signal in cells.

b. Quantification of the scaled corrected total cell fluorescence of equal size areas in the primary nucleus (PN) and micronucleus (MN) of CHP-212 cells (n=12 cells with micronuclei, two way ANOVA). Each dot represents one area.

c-d. Percentage of cells with micronuclei in 2 ecDNA containing cell lines (c: Tr-14, d: CHP-212) in the presence of different concentrations of dNTPs and at different concentrations of HU after 3 days. Each dot represents the average per experiment (n= 100-221 cells per experiment).

e. Percentage of cells with micronuclei in the isogenic glioblastoma cell line pair GBM39HSR (contains *EGFR*vIII on HSR, left) and GBM39ec (contains *EGFR*vIII on ecDNA, right) after different lengths of HU treatment (80μM) (n= 3-5, Two-way ANOVA). Each dot indicates the average per replicate.

f. Percentage of cells with micronuclei stratified by the presence of either centromere 2 or the respective amplicon in cell lines with ecDNA(ecDNA: COLO 320DM, CHP-212, STA-NB10DM, Tr-14, UKF-NB6, Lan5, SMS-KAN, KP-N-YN) or without (HSR: COLO 320HSR, NGP, IMR5/75, n=3) and by the presence or absence of HU (80μM, 72h), as quantified by FISH for centromere 2 or the respective amplicon (Two-way ANOVA). Each dot represents the average per cell line.

g. Exemplary photomicrographs of fixed cells with HSR, low ecDNA content or high ecDNA content and micronuclei (white arrowhead) in the presence of HU (80 μM, 72h).

h. Fraction of cells with micronuclei with low (bottom 30%) and high (top 30%) ecDNA content after HU treatment (COLO 320DM, CHP-212, STA-NB10DM, Tr-14, UKF-NB6, Lan5, SMS-KAN, n=1 experimental replicate, n= 98-521 cells per replicate, Paired Student's t-test). Each paired dot represents one ecDNA cell line.

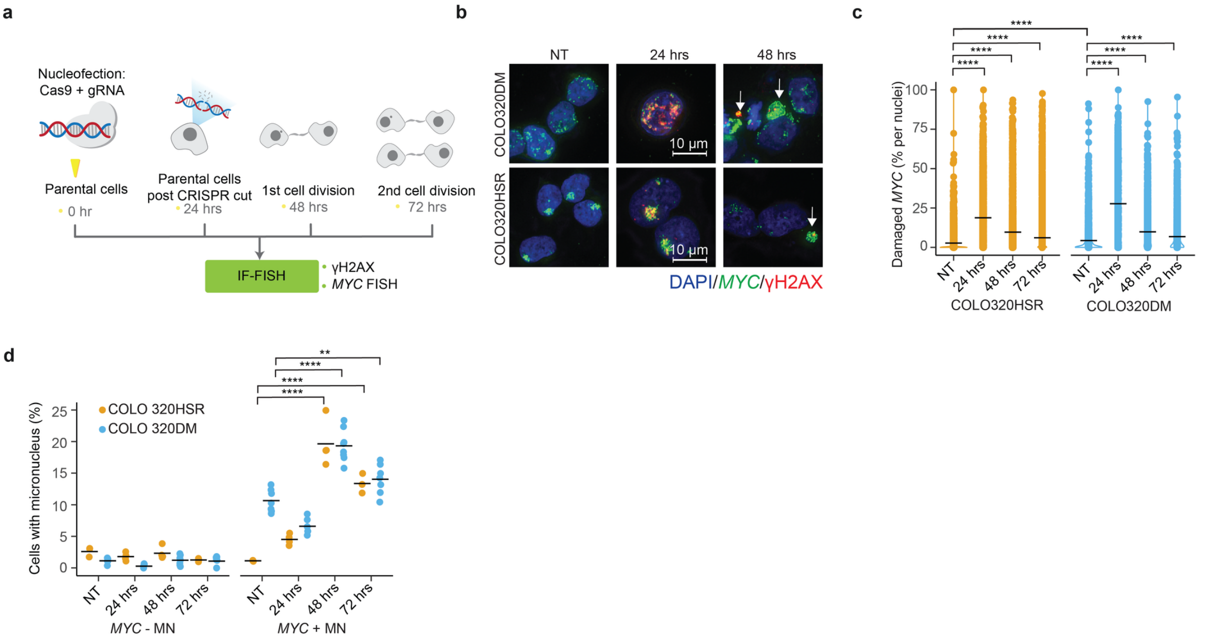


**Extended Fig. 7: Nucleofection with Cas9 induces DNA damage and micronucleation**

a. Schematics to depict samples collected at different time points.

b. Exemplary photomicrographs in COLO 320DM/COLO 320HSR cells to show DNA damage induced at amplicon$^{+}$ micronuclei formation. White arrows indicate micronuclei. Scale bar: 10µm.

c. Quantification of damaged *MYC* percentage in COLO 320DM and COLO 320HSR at different timepoints after nucleofection of Cas9+ sgRNA. (n= 2741-6696, one-way ANOVA). Each dot denotes one cell.

d. Percentage of cells with *MYC*$^{+}$ or *MYC*$^{-}$ micronuclei at different timepoints after nucleofection with guides targeting the amplicon in the isogenic cell line pair COLO 320HSR (contains *MYC* on HSR, orange) and COLO 320DM (contains *MYC* on ecDNA, blue) (n= 1919-4441, One-way ANOVA). Each dot represents one image.

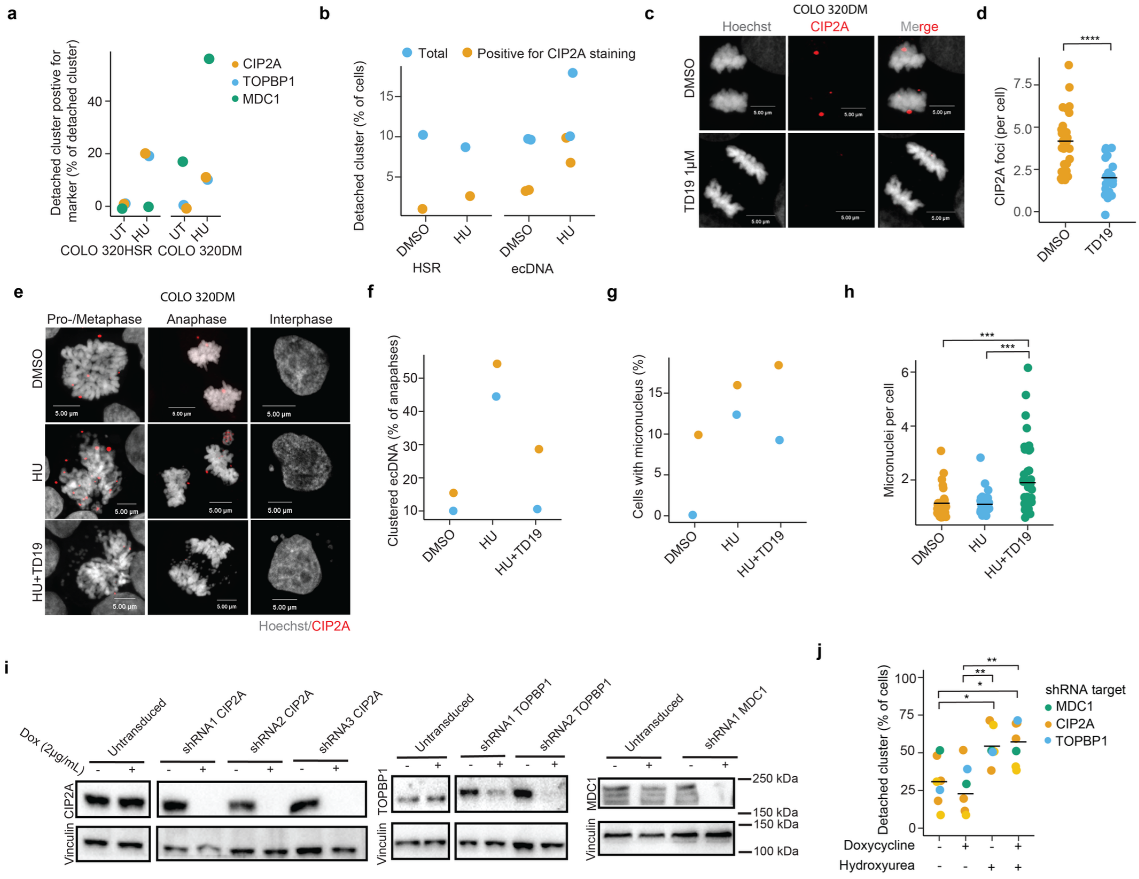


**Extended Fig. 8: Perturbation of the CIP2A-TOPBP1 complex reduces clustering of detached ecDNA**

a. Quantification of the number of detached clusters stained positive for CIP2A, TOPBP1 or MDC1 in COLO 320DM and COLO 320HSR. In the presence (HU) or absence (UT) of hydroxyurea (80μM, 24h). (n=10-12 anaphases). Each dot represents the mean per staining and treatment.

b. Percentage of detached clusters and CIP2A-positive detached clusters during anaphase in the presence (HU) or absence (DMSO) of HU (80μM, 24h). (HSR: COLO 320HSR, ecDNA: COLO 320DM and STA-NB10DM). Dots (yellow and blue) represent one individual experiment.

c. Exemplary photomicrographs of a fixed cell with ecDNA during anaphase treated with and without the CIP2A inhibitor TD19 (1 μM), stained with Hoechst (grey) and CIP2A immunofluorescence (red). Scale bar: 5μm.

d. Quantification of CIP2A foci in COLO 320DM cells during anaphase without (DMSO) and with the CIP2A inhibitor TD19 (1 μM) (DMSO, n=31; TD19, n=24; Student's t-test). Each dot represents one cell.

e. Exemplary photomicrographs of fixed COLO 320DM cells during pro-/metaphase, anaphase and interphase after treatment with DMSO, HU (80 μM) and HU (80 μM) and TD19 (1 μM) for 24h, stained for Hoechst (grey) and CIP2A immunofluorescence (red). Scale bar: 5 μm.

f. Percentage of lagging clustered ecDNA during anaphase in two ecDNA-positive cell lines (COLO 320DM, STA-NB10DM) after treatment with DMSO, HU (80 μM) or HU (80 μM) and TD19 (1 μM) for 24h. Each dots and colour represent one cell line (n= 10 anaphases per cell line and treatment).

g. Percentage of micronuclei in two ecDNA-positive cell lines (COLO 320DM, STA-NB10DM) after treatment with DMSO, HU (80 μM) or HU (80 μM) and TD19 (1 μM) for 24h. Each dot and colour represents one cell line (n= 10 anaphases per cell line and treatment).

h. Number of micronuclei per cell in COLO 320DM cells after treatment with DMSO, HU (80 μM) or HU (80 μM) and TD19 (1 μM) for 24h. (DMSO, n= 43; HU, n=43; HU+TD19, n= 40; One-way ANOVA). Line indicates the mean.

i. Western blot validation of the inducible knockdown of CIP2A, TOPBP1 and MDC1 after 2 days of 2 μg/mL doxycycline treatment in COLO 320DM after transduction with different shRNA constructs.

j. Quantification of detached clusters in COLO 320DM cells after shRNA-mediated inducible knockdown of CIP2A, TOPBP1 and MDC1 with or without the addition of doxycycline and HU (Dox: 2 μg/mL, 2d; HU: 80μM,1d).

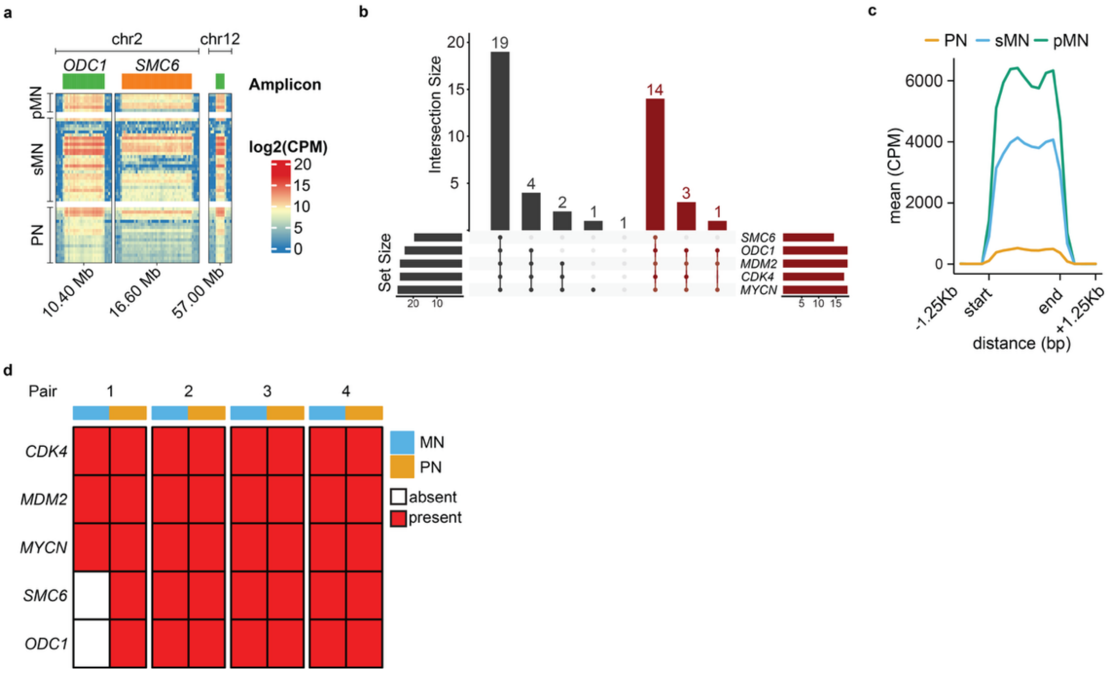


**Extended Fig. 9: Single micronucleus sequencing reveals collective entrapment of different ecDNA species**.
a. Laser microdissection directed sequencing of primary nuclei (PN) and micronuclei (MN) after incubation of TR-14 cells, containing 5 different ecDNA species, in the presence of HU (80 µM, 72h). Counts per million (CPM), normalized and log2 transformed read coverage (25kb bins) across ecDNA (*ODC1,* green; *SMC6,* orange*)* and their flanking regions (100 kb; primary nucleus, PN, n = 18; single micronucleus, sMN, n = 27; pooled micronuclei, pMN, n = 6).
b. UpSet plot of the co-occurrence of 5 ecDNA species in single micronuclei (black) and primary nuclei (red).
c. Signal plot of the mean coverage over length-scaled ecDNA in single primary nuclei (PN, n = 18, orange), single micronuclei (sMN, n = 27, blue) and pooled micronuclei (pMN, n = 6, green). Counts per million (CPM) across all ecDNA aligned and length normalized.
d. Co-occurrence of 5 ecDNA species in 4 pairs of collected micronuclei (MN) and their corresponding primary nucleus (PN) in the same cell (n=4).

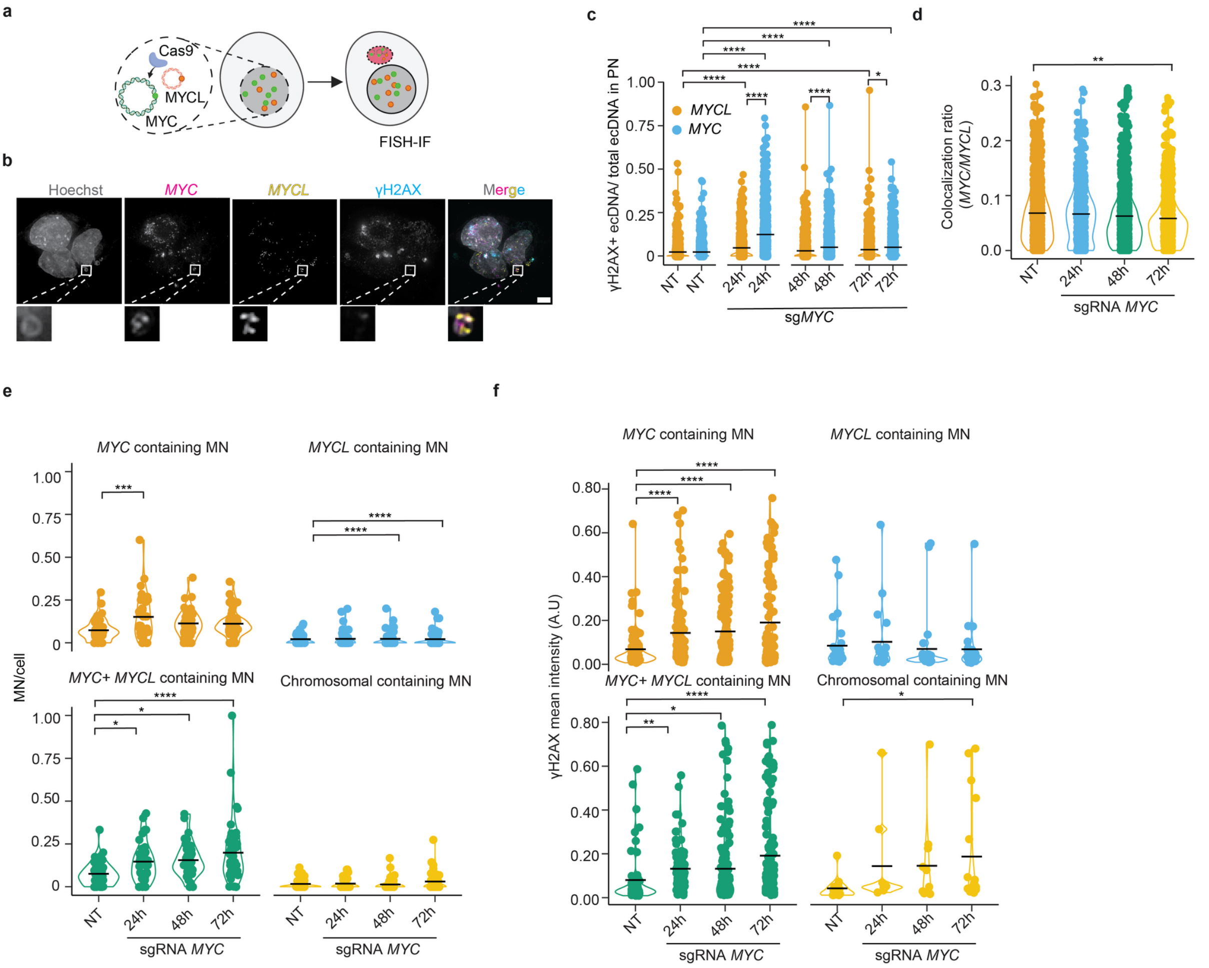


**Extended Fig. 10: Cutting of one ecDNA species induces co-micronucleation of the other ecDNA species**

a**.** Schematic of experimental design. DMS273 cells, containing *MYC* and *MYCL* on different ecDNA species are subjected to CRISPR-Cas9 mediated cutting of the *MYC* ecDNA. FISH for *MYC* and *MYCL* combined with immunofluorescence against γH2AX in fixed cells at different timepoints was used to measure the colocalisation of DNA damage and *MYC* and *MYCL* ecDNA..Scale bar: 10µm.

b. Exemplary photomicrograph of fixed DMS273 cells, containing *MYC* and *MYCL* on different ecDNA species, after sg*MYC* transfection, stained for Hoechst (grey), *MYC* FISH (magenta), *MYCL* FISH (yellow) and γH2AX immunofluorescence (blue).

c. Quantification of the fraction of γH2AX$^{+}$ *MYC* ecDNA (blue) or γH2AX$^{+}$*MYCL* ecDNA (orange) in DSM273 cells after *MYC* sgRNA transfection at different timepoints, as measured by *MYC*, *MYCL* FISH and γH2AX immunofluorescence (n= 655-1092, One-way ANOVA). Each dot represents one cell.

d. Quantification of colocalisation of *MYCL and MYC* ecDNA in fixed DMS273 cells after *MYC* sgRNA transfection at different timepoints, measured by *MYC* and *MYCL* FISH (n= 656-1091, One-way ANOVA). Each dot represents one cell.

e. Fraction of micronuclei, normalized by cell number, containing *MYC* ecDNA, *MYCL* ecDNA, *MYC* and *MYCL* ecDNA or other chromosomes in fixed DSM273 cells after *MYC* sgRNA transfection, quantified by *MYC* and *MYCL* FISH (n=38-54, One-way ANOVA). Each dot represents one image.

f. Quantification of γH2AX mean intensity in micronuclei in fixed DMS273 cells after *MYC* sgRNA transfection stratified by micronucleus content, measured by *MYC*, *MYCL* FISH and γH2AX immunofluorescence (n= 92-128, One-way ANOVA). Each dot represents one micronucleus.

**a**

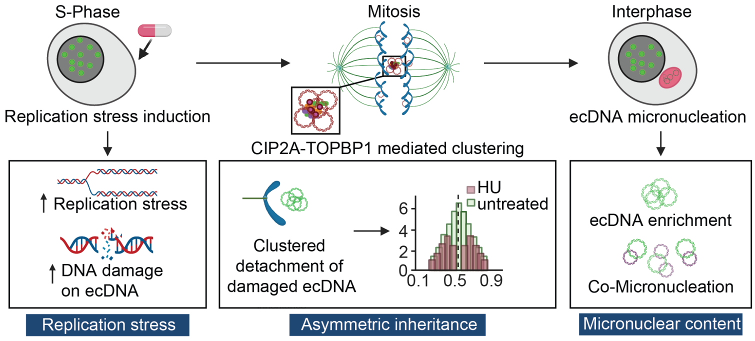


**Extended Fig. 11: Schematic model illustrating the pathway from replication stress induction to the formation of ecDNA-containing micronuclei.**

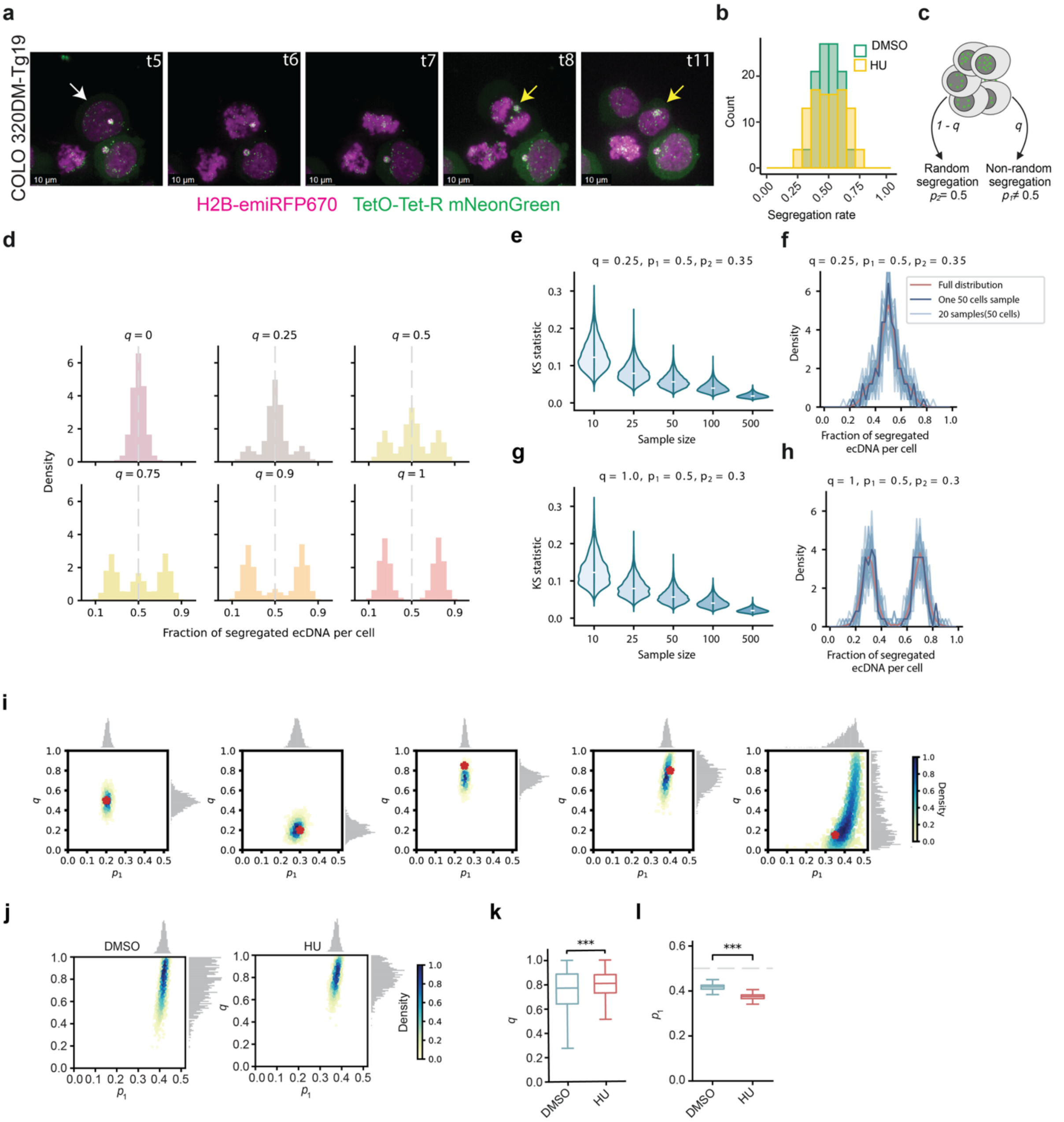


**Extended Fig. 12: Asymmetric inheritance of ecDNA⁺ MN by live cell imaging and a biased random-segregation model.**

a. Live-cell imaging of the COLO 320DM-Tg19 cell line engineered to visualize *MYC* ecDNA (TetO-TetR-mNeonGreen) during mitotic transition. Representative snapshots of a mitotic cell (white arrow) carrying an ecDNA-containing micronucleus (yellow arrow) getting asymmetrically inherited into one of the daughter cells upon mitotic exit. Scale bar: 10μm.

b. Histograms of ecDNA fractions in the CHP212 cell line in DMSO treated cells without visible lagging clusters (left), DMSO treated cells (middle) and HU treated cells (HU), showing the distribution of ecDNA segregation across daughter cells (n= 50 anaphases untreated, n= 50 anaphases after 24h HU treatment).

c. Overview and simulation results of the biased random segregation model.

d. Simulated distributions of ecDNA fractions under varying $q$ values (proportion of biased segregation), with $p_1 = 0.25$ (extent of biased segregation).

e-h. Comparison of ecDNA fraction distributions under different sample sizes. (e) Violin plot showing the Kolmogorov distances between the sampled distributions and the full distribution across varying sample sizes. (f) Comparison of ecDNA fraction distributions: the red line represents the full distribution generated from $10^7$ cell divisions, while the blue line represents the distribution based on 50 sampled ecDNA fractions. (g, h) Same analyses as shown in (e) and (f), respectively, but generated using different parameters as indicated in the figure.

i. Density scatter plot of inferred parameters $p_1$ and $q$ using synthetic data (50 pairs of daughter cells) for different combinations $p_1$ and $q$. The marginal distributions of $p_1$ and $q$ are displayed above and to the right, respectively. Density values are normalized to 1. The red star indicates the true parameter values.

j. Density scatter plot of inferred parameters $p_1$ (extent of biased segregation) and $q$ (proportion of biased segregation), with untreated (DMSO) cells on the left and HU-treated cells on the right. The marginal distributions of $p_1$ and $q$ are shown above and to the right, respectively. Density values are normalized to 1.

k. The inferred proportion of cells exhibiting biased segregation is significantly increased in drug-treated cells compared to untreated cells. Statistical significance was determined using the Mann–Whitney U test ($p < 0.001$).

l. The inferred $p_1$ is significantly reduced in drug-treated cells compared to untreated cells. Statistical significance was determined using the Mann–Whitney U test ($p < 0.001$).

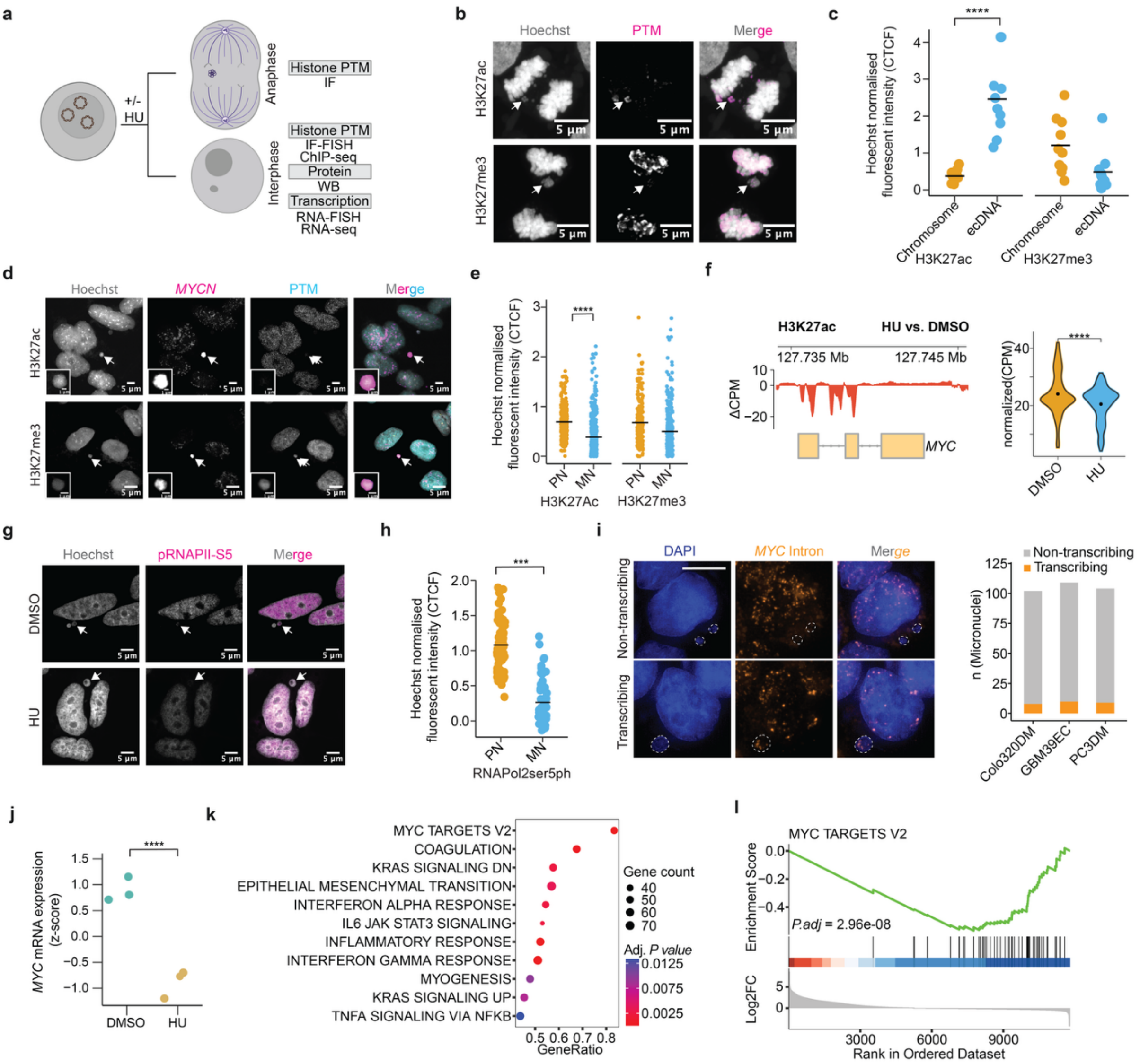


**Extended Fig. 13 Reduced oncogenic transcription of ecDNA in micronuclei.**

a. Schematic of experimental setup

b. Exemplary photomicrographs of COLO 320DM cells with *MYC* ecDNA in anaphase stained using immunofluorescence against H3K27Ac or H3K27me3 (ecDNA is indicated with white arrowheads). Scale bar: 5 µm

c. Ratio of H3K27Ac and H3K27me3 intensity on COLO 320DM *MYC* ecDNA vs. linear chromosomes (n=9). Each dot represents one detached ecDNA or linear cluster in anaphase.

d. Exemplary photomicrographs of a Tr-14 cells with *MYCN* ecDNA stained using immunofluorescence against H3K27Ac or H3K27me3 (micronuclei are indicated with arrowheads). Scale bar: 5 µm

e. Hoechst normalized H3K27Ac and H3K27me3 fluorescence intensity in micronuclei vs. primary nuclei, quantified by immunofluorescent staining in Tr-14 and CHP-212 (n= 221- 470). Each dot represents one primary or micronucleus.

f. ChIP-sequencing of H3K27ac in COLO320DM cells. Left panel: from top to bottom: peak call annotation. DMSO-subtracted read density as counts per million (CPM). Gene annotation. Right panel: Input-subtracted read density as CPM.

g. Exemplary photomicrographs of Tr-14 cells with *MYCN* ecDNA incubated with hydroxyurea (80 µM) or DMSO (vehicle control) and stained using immunofluorescence against RNA polymerase II phosphorylation at serine 5 (micronuclei are indicated with arrowheads).

h. Hoechst normalized intensity in micronuclei vs. primary nuclei from Tr-14 cells stained using immunofluorescence against RNA polymerase II phosphorylation at serine 5 (n=32-35). Each dot represents one primary or micronucleus.

i. Exemplary photomicrographs of COLO320DM cells and their micronuclei (white dashed line) labelled with intron *MYC* RNA FISH signal (left), showing a non-transcribing micronucleus (top) and an actively transcribing micronucleus (bottom). A quantification of the number of transcribing and non-transcribing micronuclei in COLO320DM (n=102), GBM39EC (n=109), and PC3DM (n=104) (right). Scale bar: 10µm.

j. Z-score normalized transcript counts of the circularly amplified *MYC* oncogene in COLO320DM cells. Three replicates per treatment condition. Benjamini-Hochberg procedure-corrected p-value is shown (Wald test).

k. Significantly enriched MSigDBhallmark genesets between hydroxyurea and DMSO treated cells. Benjamini-Hochberg procedure corrected p-values and gene counts are shown.

l. Gene set enrichment analysis (GSEA) of MYC target genes. Benjamini-Hochberg procedure-corrected p-value is shown.

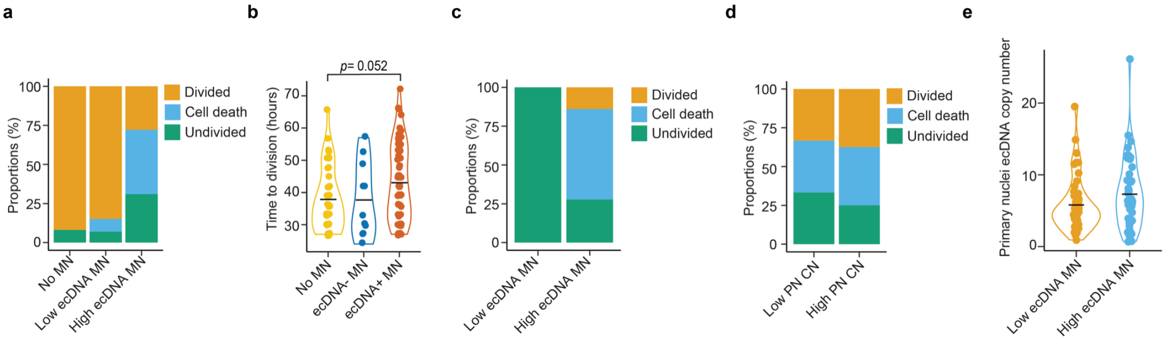


**Extended Fig. 14: Fitness in cells with $ecDNA^+$ MN is correlated with micronucleus size, but not the ecDNA copy number in primary nucleus**

a. Quantification of COLO 320DM-Tg19 cell fates as determined using live cell imaging in the absence of HU for cells without micronuclei (No MN, n=17), with MN containing low ecDNA content (n=18), or with MN containing high ecDNA content (n=13). Cell fates include cell division, cell death, or remaining undivided.

b. Time to the next cell division determined by live cell imaging in COLO 320 DM-Tg19 cells pre-treated with hydroxyurea for 24 hours (no MN n= 49, $ecDNA^-$ MN n=13, $ecDNA^+$ MN n= 58). Data is collected from three independent experiments.

c. Fate of daughter cells bearing micronuclei of low or high ecDNA content, as determined by whether the ecDNA content in MN is below (low) or above (high) the median of ecDNA content in all MN over 2 µm in diameter (Low ecDNA MN n=2, High ecDNA MN n=36). Outcomes include continued division, cell death, or remaining undivided at the experimental end point. Data is collected from three independent experiments.

d. Fate of daughter cells stratified by low or high ecDNA content in PN, as determined by whether the ecDNA content in PN is below (low, n=55) or above (high, n=37) the mean of ecDNA content in all PN. Outcomes include continued division, cell death, or remaining undivided. Each bar plot depicts an individual experiment.

e. Copy number of ecDNA in the primary nuclei in cells with low (below median, n=46) or high (above median, n= 46) ecDNA content in micronuclei.

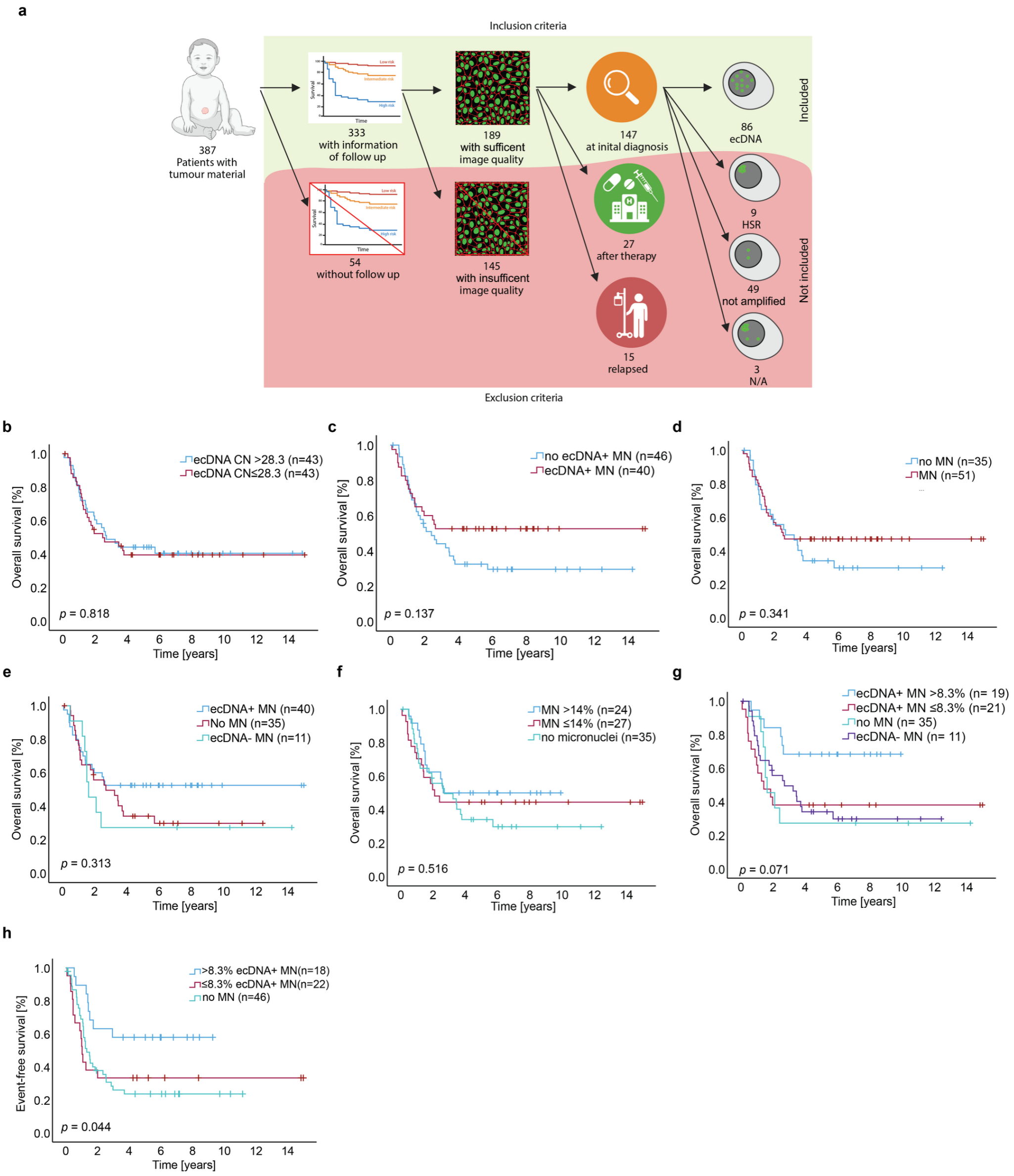


**Extended Fig. 15: High ecDNA micronucleation improves patient survival**
a. Schematic showing the inclusion and exclusion criteria for patient selection in the following analysis.
b. Kaplan–Meier survival curves for patients with above or below median copy number (CN) in primary nucleus determined by FISH (OS, log-rank test).
c. Kaplan–Meier survival curves for patients with or without $ecDNA^+$ MN, as determined by FISH signal in the MN (OS, log-rank test).
d. Kaplan–Meier survival curves for patients with or without MN (OS, log-rank test).
e. Kaplan–Meier survival curves for patients with $ecDNA^+$ MN, $ecDNA^-$ MN or without MN, as determined by FISH signal in the MN (OS, log-rank test).
f. Kaplan–Meier survival curves for patients with above or below median MN frequency or no MN (OS, log-rank test).
g. Kaplan–Meier survival curves for patients with >8.3% $ecDNA^+$ MN, <8.3% $ecDNA^+$ MN, ecDNA- MN or no MN, as determined by FISH signal in the MN (OS, log-rank test).
h. Kaplan–Meier survival curves for patients with >8.3% $ecDNA^+$ MN, <8.3% $ecDNA^+$ MN, or no MN, as determined by FISH signal in the MN (Event-free survival, log-rank test).
